# Wings expansion in *Drosophila melanogaster*

**DOI:** 10.1101/2024.05.17.594352

**Authors:** Simon Hadjaje, Ignacio Andrade-Silva, Marie-Julie Dalbe, Raphaël Clément, Joel Marthelot

## Abstract

During their final transformation, insects emerge from the pupal case and deploy their wings within minutes. The wings deploy from a compact origami structure, to form a planar, rigid and functional blade that allows the insect to fly. The deployment is powered by a rapid increase in internal pressure, and by the subsequent flow of hemolymph into the deployable wing structure. Using a combination of imaging techniques, we characterize the internal and external structure of the wing in *Drosophila melanogaster*, the unfolding kinematics at the organ scale, and the hemolymph flow during deployment. We find that beyond the mere unfolding of the macroscopic folds, wing deployment also involves an expansion of cell surface and the unfolding of microscopic wrinkles in the cuticle enveloping the wing. A quantitative computational model, incorporating mechanical measurements of the viscoelastic properties and microstructure of the wing, predicts the existence of an operating point for deployment and captures the dynamics of expansion. This model suggests that insects exploit material and geometric nonlinearities to achieve rapid and efficient reconfiguration of soft deployable structures.

---

Embryonic development is the stage of major morphological transformations of tissues and organs [1±5]. The biological and physical mechanisms by which these transformations are coordinated in space and time are a central issue in developmental biology. The wing of the fruit fly *Drosophila melanogaster* is a paradigmatic example of such continuous organ-scale transformation, combining expansion and buckling of the wing epithelium to shape the pupal wings into a highly folded origami structure [6±9]. However, how the origami unfolds into a functional wing within minutes after emergence remains largely unknown. Early work has suggested a mechanical basis for wing expansion in insects, associated with an increase in hemolymph pressure in the open circulatory system 10±12] under hormonal control [13, 14], but an integrative physical model of this transformation is lacking. Here, we describe wing actuation as a fluid-structure interaction problem based on experimental characterization of mechanical properties and unfolding kinematics at organ and tissue scales. We show that wing pressurization, combined with constitutive properties and tissue geometry, results in an operating point that the insect uses for robust unfolding. This work sheds light on a neglected but fascinating morphogenetic process at the organ scale. It will also inform engineering strategies for deployable structures whose shape evolves from compact and folded to extended and operational, with applications ranging from biomedical to aerospace [15], and for morphing artificial matter that continuously changes shapes [16], with applications ranging from programmable mechanical metamaterials [17±21] to soft robotics [22±25].

## Hemolymph injection into a two-dimensional network causes wing expansion

We first design a protocol to image wing deployment macroscopically. Upon eclosion, the fly is glued to a thin fiber mounted on a micromanipulator, and expansion is recorded using bright-field microscopy from dorsal and lateral views. Fig.1a shows snapshots of the dorsal view of the wing expansion (see Supplementary Movie 1). The wings start out highly folded, with macroscopic folds along the longitudinal veins. The two wings expand simultaneously, dramatically changing shape from a compact three-dimensional structure to a fully deployed flat wing (Fig.1a). Additional characteristic features of deployment are revealed by lateral observation. The strong activity of the pharyngeal pump (see Supplementary Movie 2) indicates that the fly swallows air prior to wing expansion [26], resulting in inflation of the ptilinum, a reversible pouch located on the fly’s head. The abdominal contractions that occur simultaneously contribute to an increase in internal pressure, which results in the injection of hemolymph into the wing, causing it to expand [10]. At the end of expansion, the ptilinum deflates and the abdomen relaxes, indicating a decrease in internal pressure. After expansion, the final shape of the wing is stabilized by cuticle tanning and sclerotization [27].

To gain insight into the microscopic structure of the wing, we used high-resolution X-ray micro-CT just before deployment (Fig. 1b(i) and Supplementary Movie 3). A cross-section normal to the proximo-distal axis (Fig. 1b(ii)) reveals that the wing is composed of two continuous, folded plates of thickness *e* = 6.5 *μ*m, connected by regularly spaced pillars of height *h* = 7.5 *μ*m. Veins, hollow tubular structures indicated by white arrows on Fig. 1b(ii), are already visible in the folded wings. The cross-section normal to the dorso-ventral plane reveals that the pillars are arranged in a regular hexagonal lattice of period *a* = 6.7 *μ*m and diameter *d* = 3.3 *μ*m (Fig. 1b(iii)). The inside of the wing thus forms a two-dimensional porous network as sketched in Fig. 1b.

**Figure 1:**
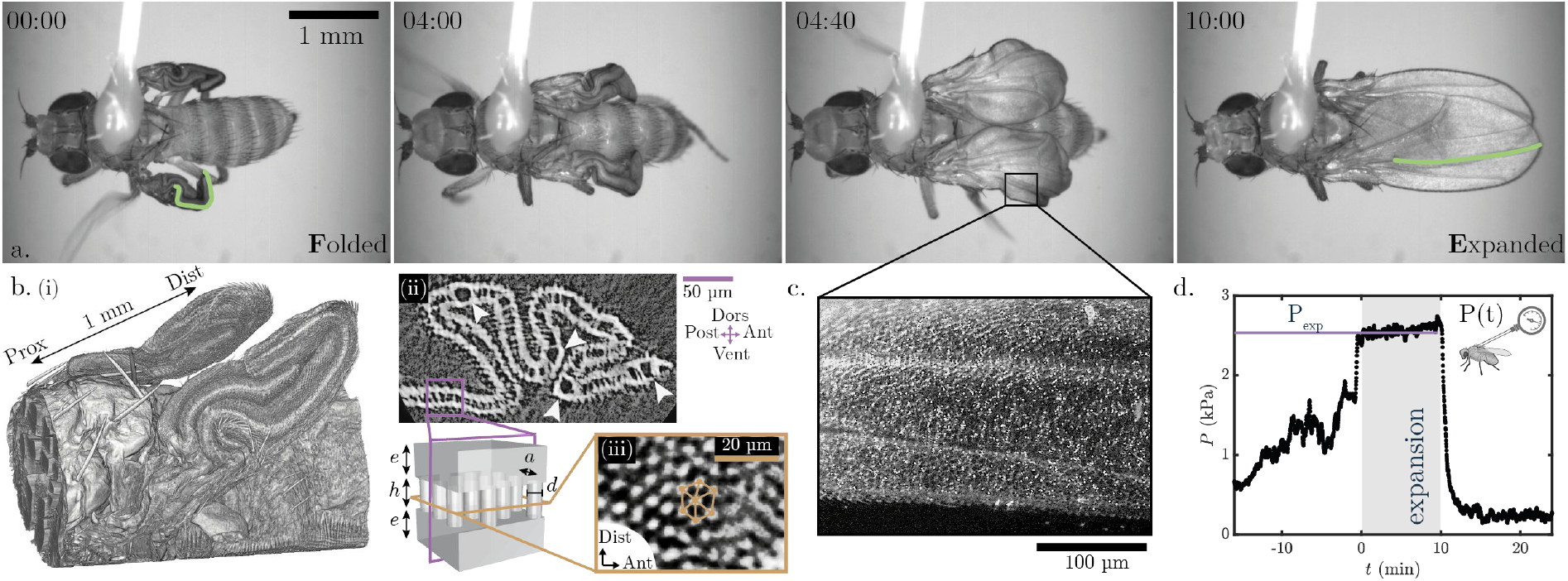
Geometry and pressure in the unfolding wing. (**a**) Snapshots of the wings expansion with one of the longitudinal vein highlighted in green. **(b)** (i) Microtomography of the folded wings. (ii) Micro-CT scan cross-section showing macroscopic folds, vein structure (white arrows) and internal pillars. (iii) Perpendicular section revealing the hexagonal pillars organization. The sketch summarizes the wing structure: two plates (thickness *e*) connected with pillars (height *h*, diameter *d*) organized in an hexagonal lattice (interpillar distance *a*). **(c)** Fluorescent beads (white dots, hemolymph markers) flowing in an expanding wing. **(d)** *in vivo* recording of the internal pressure *P* (*t*) of a newly emerged insect. Wings expansion takes place throughout the grayed-out segment at a constant pressure *P*_*exp*_.

To understand how the rise in pressure guides the structure deployment, we first examine the hemolymph distribution in the wing. Prior to expansion, fluorescent beads are injected into the abdomen of the fly. Fig. 1c shows that the beads permeate the entire wing surface during expansion, indicating that hemolymph is injected throughout the structure in the two-dimensional porous network, rather than just confined to the veins as observed in adults (see [28] and Supplementary Movie 4), and commonly assumed during deployment [7, 29].

The internal pressure during expansion is measured by stinging the thoracic region between the two wings (the scutellum) of newly emerged flies, with a capillary connected to a pressure transducer (see Supplementary Movie 5). The pressure increases over approximately 15 minutes before reaching a plateau of *P*_*exp*_ ≈ 2.5 kPa, as shown in Fig. 1d. The wings expand at this constant pressure. Following expansion, the pressure suddenly drops to a lower value of a few hundred pascals. This coincides with the rapid deflation of the ptilinum, that is highly inflated during expansion, and the relaxation of muscle contractions. The abdomen goes from an elongated and stretched state to a shorter and relaxed state. The abdominal and thoracic muscles involved in the increase in pressure degenerate shortly after deployment [30]. The fly thus transiently generates a few kilopascals to deploy its wings within a few minutes. Remarkably, this transformation bears similarity to that of artificial inflatable structures, such as air mattresses or paddle boards, in which pressure guides in-plane expansion of two plates connected by pillars [31].

## The wing not only unfolds but also stretches

To quantify expansion, we use the natural landmarks provided by the wing veins. The three-dimensional position of the vein network is reconstructed using micro-CT images in the folded state (Fig. 1b(iii)). This network is preserved in the deployed wing, allowing for a direct comparison between the folded and expanded states (Fig. 2a and Fig. S1). To characterize the kinematics, we first measure the apparent wing stretch Λ = *L/L*_*F*_, where *L* is the length of the segment joining the extremities of a longitudinal vein, and *L*_*F*_ the corresponding initial folded length as shown in the inset in Fig. 2b. The apparent stretch Λ is thus dominated by the kinematic opening of macroscopic folds. We find that expansion dynamics are highly reproducible between individuals (see Supplementary Movie 6), with wing overall length increasing by a factor of 3 within 2 minutes (Fig. 2b).

**Figure 2.**
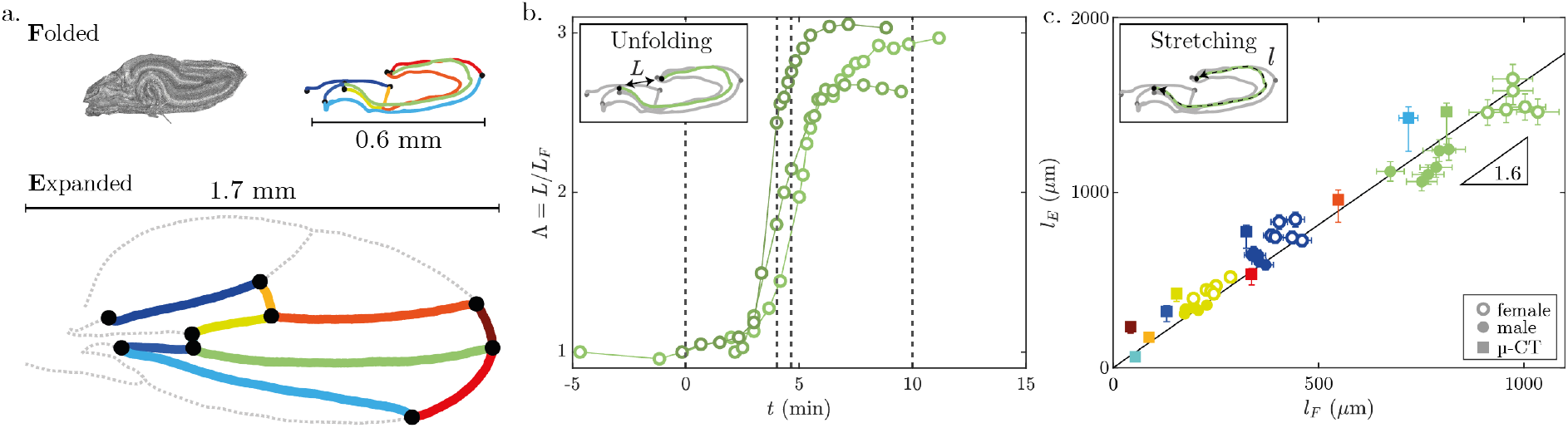
Wing elongation during expansion. **(a)** Veins network in folded state (from micro-CT scans) and adult state (from bright field microscopy images). Each color represents a specific vein. **(b)** Apparent wing stretch Λ = *L/L*_*F*_ as a function of time for 3 different flies. *L* is the length between the two ends of the longitudinal vein highlighted in green in (a) and Fig. 1a, and *L*_*F*_ the initial folded length. Dotted vertical lines correspond to the snapshots in Fig. 1a. **(c)** Arclength of expanded veins *l*_*E*_ versus folded veins *l*_*F*_. Each color represents a vein shown in (a). Length measurements from micro-CT are represented by square markers, while veins tracked on videos are represented by circular markers (empty for females, full for males). Solid line is a linear fit of all experiments. Standard deviation and relative uncertainties due to video tracking are indicated by error bars.

Next, we investigate whether the deployment of the folded wing is solely due to the kinematic opening of macroscopic folds, similar to the unfolding of inextensible origami paper [17±19] or the wing of adult Coccinellidae [32], or whether it involves a combination of structural unfolding and tissue stretching. To measure changes in tissue metrics, we compare the arclength of veins in the folded and expanded states. In Fig. 2c, we plot the arclength of the expanded vein *l*_*E*_ against the initial folded arclength *l*_*F*_. To complement these measurements, we track deformation of three individual veins (marked in blue, yellow, and green in Fig. 2a) using bright-field microscopy observations of dorsal and lateral views (circular markers in Fig. 2c). Notably, we find that the wings not only unfold during expansion, but also lengthen by approximately 1.6 times their original length. Veins oriented in different directions present the same stretch, indicating that extension is isotropic in the plane. Therefore, the rapid and dramatic expansion of the wings appears to be a dual morphing response that involves unfolding macroscopic folds and additional isotropic tissue stretching. Next, we address how this tissue stretching is manifested at the cellular level.

## Apparent stretching relies on unwrinkling at the cellular scale

In order to gain a deeper understanding of the kinematics of unfolding at the cellular level, we conduct observations of cross-sections of wings normal to the proximo-distal axis (dotted plane in the sketch in Fig. 3a) using transmission electron microscopy (TEM) in both the folded and unfolded states. Micrographs (Fig. 3b) show that the folded wing is composed of two layers of epithelial cells (one ventral, the other dorsal, shown in green) separated by a lumen, into which hemolymph flows. The epithelial cell layer is covered by a thin, wrinkled layer of cuticle (shown in red). During the process of expansion, the cuticle unwrinkles while maintaining a constant thickness of 200 nm in the adult wing. This suggests that the cuticle flattens without undergoing any significant stretching. The corresponding micrograph (Fig. 3b, bottom) is obtained a few hours after the expansion process, at which point the epithelial cells have already undergone apoptosis, detached, and been washed out of the wing by the action of the wing hearts [33, 34], leaving only the cells surrounding the veins. The variation in wing thickness under pressure is measured using two-photon microscopy. We observe that the pillars stretch under hemolymph pressure, increasing the space between the two cell layers during expansion (see wing thickness in Fig. 3e, Fig. S2d and Supplementary Movie 7).

**Figure 3.**
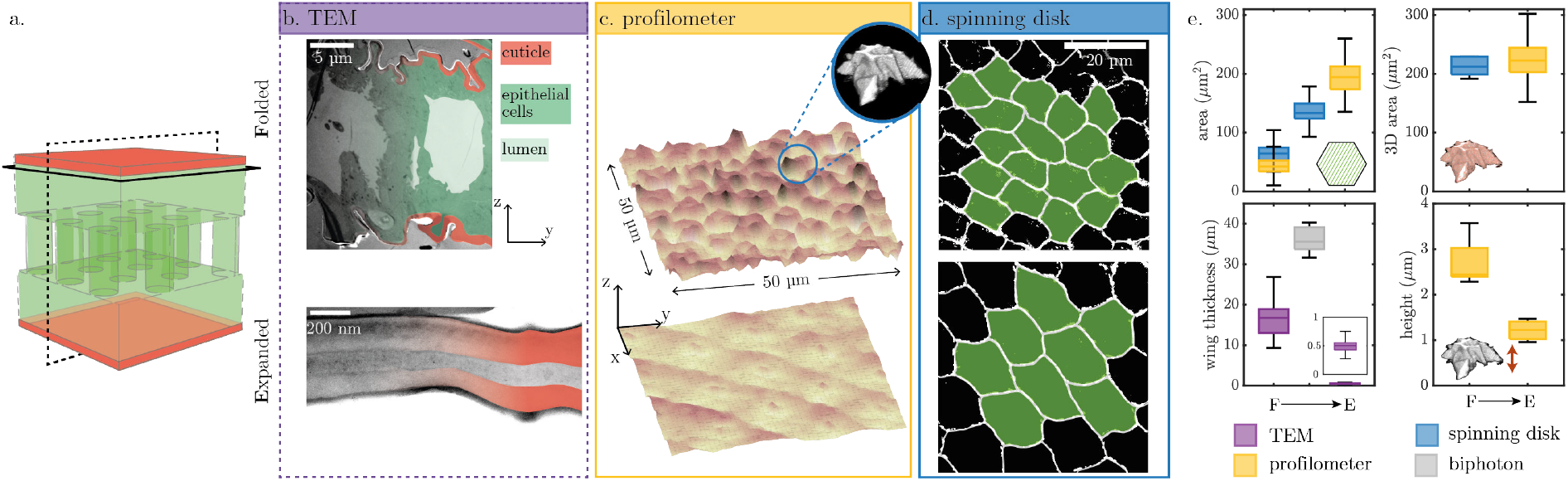
Cellular scale unwrinkling. **(a)** Schematic view of a wing microscopic structure: 2 monolayers of epithelial cells (green) covered by a rigid cuticle (red) and connected by pillars. **(b)** Cross section following the dotted frame in (a), transmitted electron microscopy (TEM). **(c)** Wing surface scans (optical profilometer, 50×50 *μ*m^2^). Top right inset: 3D reconstruction of the apical surface of an epithelial cell (spining disk confocal microscopy, Utrophin:GFP). **(d)** Epithelial cells contour (spining disk confocal microscopy, Ecad:GFP). For (b-d) top images correspond to folded wings, while bottom images correspond to expanded wings. **(e)** Evolution of the main geometric characteristics from folded (F) to expanded (E) stages (from top left to bottom right diagram): 2D epithelial cell surface area; 3D surface area integration of the cuticle; total wing thickness; and wrinkles height. Data are obtained by TEM (purple boxes, see (b)), profilometer scans (yellow, (c)), spinning disk (blue, (d)) and biphoton images (gray, see Fig. S2d in supplementary).

Using optical profilometry, we measure the topography of the apical surface of the folded and expanded wing (solid line plane in the sketch in Fig. 3a). The wrinkles are organized in a hexagonal pattern, corresponding to the cellular tiling (see Fig. 3c). From cell centroids, we generate the Voronoi tessellation in order to compute the 2D surface area and maximum height of the wrinkles. Fig. 3e shows that the height of the wrinkles decreases while the 2D surface area increases during expansion, indicating a general flattening of the wing surface. This increase of cell surface area is confirmed by direct measurement in the folded state and during expansion using spinning disk microscopy (Fig. 3d). Note that the cell shape after expansion shows no significant shape anisotropy, suggesting isotropic cell stretching. We label the cells’ apical surface, i.e. the surface where the cuticle adheres, to segment individual cells and reconstruct their shape in 3D, as shown in the inset of Fig. 3c. In folded wings, apical surface of the cells is shaped like a volcano (see Supplementary Movie 8). The integration of the 3D outer surface (see boxplot 3D area in Fig. 3e) reveals that the cuticle has sufficient surface area to unwrinkle without stretching. Moreover, the volcano initial shape is isometric with respect to a plane, and allows the cuticle to unwrinkle without stretching during wing expansion.

## Coupling between constitutive properties and geometry reveals an operating point for deployment

The kinematic observation of the wing microstructure is summarized in Fig. 4a. The ventral and dorsal plates are each composed of a bilayer of epithelial cells covered by a thin, rigid, and wrinkled cuticle. During expansion, the epithelial cells flatten their apical surface, thus expanding their area, while the rigid cuticle unwrinkles without stretching. The two plates are connected by regularly spaced pillars organized in a hexagonal lattice. Large microtubule bundles extend from the apical junctions to the basal junctions of the cells [35, 36] and straighten under pressure, thereby limiting the separation of the two plates.

**Figure 4.**
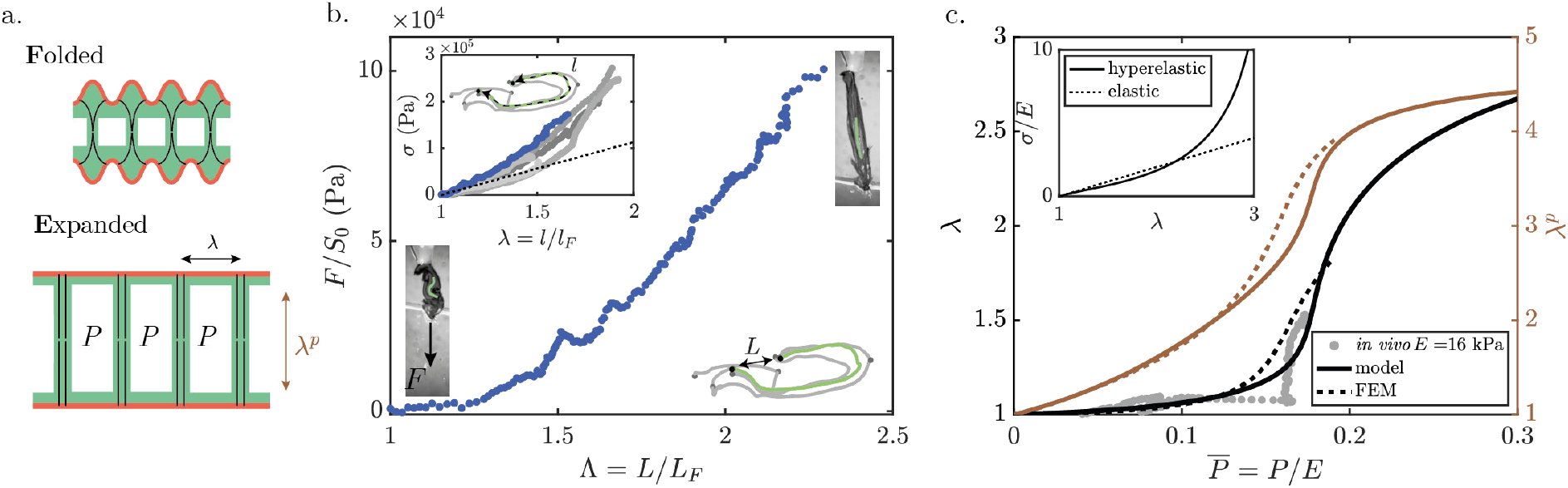
Mechanics of the wing expansion. **(a)** Cross-sectional sketch of the folded (top) and expanded (bottom) wing structure with wrinkles and microtubules under pressure. **(b)** *F/S*_0_ versus apparent stretch Λ obtained from tensile test of the wing. (Inset) true stress *σ* = *λF/S*_0_ versus stretch *λ* (linear elastic model in dotted line). **(c)** Prediction of in-plane stretch *λ* (black, left axis) and pillars vertical stretch *λ*^*p*^ (orange, right axis) as a function of the normalized applied pressure 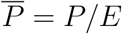 for a hyperelastic Gent model (model: solid lines; FEM: dashed lines). Pressure measurements normalized by a Young’s modulus of *E* = 16 kPa are shown as gray markers. (Inset) Constitutive law for a linear elastic model (dotted line) and a hyperelastic Gent model with *J*_*m*_ = 20 (solid line).

To gain insight into the mechanics of deployment, we start out by conducting tensile tests on dissected folded wings. One wing extremity is fixed to a motorized linear stage, and a constant velocity of 10 *μ*m/s is imposed while the other wing extremity is attached to a rigid fiber connected to a load sensor (see Supplementary Movie 9).

The force *F* is normalized by the wing’s initial cross-sectional surface area *S*_0_, which was obtained through micro-CT scanning. In Fig. 4b, we plot *F/S*_0_ as a function of the apparent wing stretch Λ = *L/L*_*F*_. Two distinct regimes are observed: (i) for small values of Λ ≤ 1.25, the macroscopic folds open and the material undergoes negligible stretching. The stress remains close to zero, showing that almost no force is required to unfold the origami-like macroscopic folds. (ii) Once the main folds open, the wing stretches and stresses increase. To disentangle the effect of macroscopic unfolding and determine the actual stiffness of the material, we follow the arclength *l* of a longitudinal vein. In the inset of Fig. 4b, we plot the true stress *σ* = *Fλ/S*_0_ assuming incompressibility as a function of stretch *λ* = *l/l*_*F*_. We extract the Young’s modulus of the material *E* ≈ 59 − 147 kPa which captures the linear relationship observed at low strain (interval from *n* = 6 experiments). Stiffening is observed at large strains, which we interpret as the effect of unwrinkling of the rigid cuticle layer as the tissue expands (see Fig. S3c).

To make sense of these observations, we construct a mechanical model that couples structural deformation to hemolymph pressure increase. Deformation involves in-plane stretching of the plates *λ* and elongation of the pillars *λ*^*p*^, as well as thinning of the plates and pillars due to volume conservation (see sketch in Fig. 4a). Stresses in the pillars and plates compensate for the increase in internal pressure *P*. To capture the large-strain stress stiffening due to the in-plane unwrinkling process and the straightening of the microtubules along the pillars, we consider a phenomenological hyperelastic Gent model (see solid line in Inset Fig. 4c) with two parameters, the Young’s modulus *E* and the stress stiffening captured by the limiting value *J*_*m*_. We find that *λ* and *λ*^*p*^ are solutions of a differential non-linear system of equations [37], which we solve numerically (see Methods for computation details). Fig. 4c presents the theoretical predictions for *λ* and *λ*^*p*^ as a function of the applied non-dimensional pressure 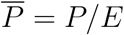 (black and orange line, respectively). The model is supported by FEM numerical simulations and predicts an operating point 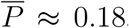, at which the wing undergoes considerable deformation while maintaining a low operating pressure. At this operating pressure, the predicted vertical stretch of the pillars *λ*^*p*^ ∼ 4 is in good agreement with the measurement of the total wing thickness obtained by two-photon microscopy (Fig. 3e). By fitting our experimental measurements on the 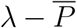 diagram, we find a Young’s modulus of 16 kPa (gray markers in Fig. 4c), which is slightly smaller than the value obtained with the tensile test.

To further evaluate the fluid-structure model, we artificially inflate fixed flies after they emerge from their pupal cases (see Methods and Supplementary Movie 10). We observe a very nonlinear behavior in plane deformation and pressure, consistent with our theoretical model. When the pressure is maintained at a plateau below 10 kPa, the wings do not expand. Expansion occurs at operating pressures between 10 and 16 kPa. At pressures above 17 kPa, the microtubules do not support pillar expansion, and we observe blisters and balloon-like wings. With the Young’s modulus measured in the tensile test, we obtain an operating point of *P/E* ∼ 0.15, which is consistent with the one predicted by the model (see Fig. 4c). While the flies are wild-type, the wings tend to curl up when deployed under artificial conditions similar to the phenotypes observed in *Curly* flies. This could indicate differential evaporation between the dorsal and ventral plates, leading to differential variation in the elastic properties of the plates. Alternatively, it may be indicative of an active control mechanism that regulates the in-plane expansion of the wings when the fly is alive.

## Expansion dynamics is governed by the viscoelastic properties of the tissue

We now seek to gain insight into the dynamics of expansion. A first hypothesis would be that the observed timescale results from the fluid-structure interaction between the flow of viscous hemolymph and the deformation of the flexible porous network. For a typical hemolymph viscosity *η*_*f*_ ∼ 1 mPa.s [38, 39], the characteristic timescale *η*_*f*_ *L*^2^*/*(*Ehe*) based on the coupling between wing elasticity and fluid viscosity [40] is on the order of milliseconds. This is several orders of magnitude shorter than the observed deployment timescale, indicating the presence of an additional dissipation mechanism.

In order to ascertain the viscoelastic properties of the wing, we perform nanoindentation measurements on the folded wing. A force *F*_*i*_ is applied at different loading rates while the indentation depth *δ* is measured (Fig. 5a). The effective stiffness appears to increase with the loading rate, a feature typical of viscoelastic materials. This indicates the existence of another source of dissipation in the material, with a characteristic time *τ* = *η/E*_*i*_ where *η* is the viscosity of the material and *E*_*i*_ the effective elastic response in indentation, with contributions from the epithelial cells as well as the cuticle.

**Figure 5.**
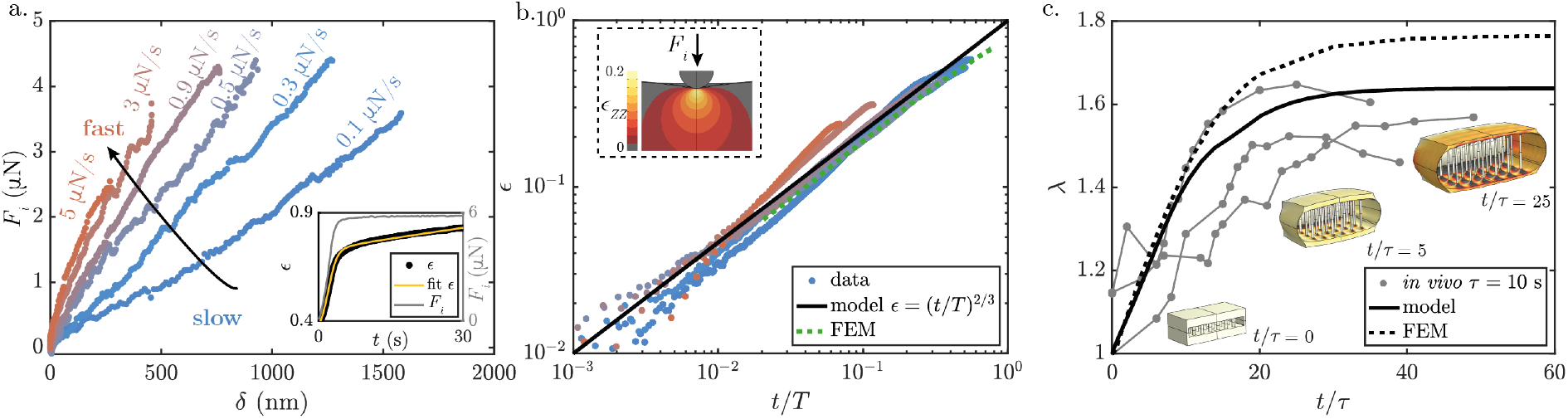
Dynamics of the wing expansion. **(a)** Force-displacement curves obtained from nano-indentation experiments on folded wings, from slow indentation (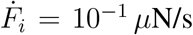, blue) to faster (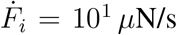, red). Inset: Deformation *ϵ* = (*δ/R*)^1*/*2^ measured over time for a creep experiment during which a constant force *F*_*i*_ is applied (gray line). Experimental data (in black) are fitted with the model (yellow line). **(b)** *ϵ* versus normalized time *t/T* : both experimental data (from (a), colored markers) and FEM numerical simulations of viscoelastic bilayer indentation (green dashed line) collapse on the model *ϵ* = (*t/T*)^2*/*3^ (black line). Inset: local strain in the indentation direction *ϵ*_*ZZ*_ from FEM simulations. **(c)** In-plane deformation *λ* as a function of *t/τ*. Experimental measurements (normalized with *τ* = 10 s): gray markers; model: solid line; FEM: dashed line. Snapshots of FEM simulations at *t/τ* = 0, 5 and 25 are shown (color bar: strain in one of the in-plane directions)

Creep experiments (inset of Fig. 5a), which consist of a loading phase at constant rate followed by a constant-force relaxation phase, are accurately represented by a three elements Maxwell-Jeffrey model comprising a dash-pot in series with a dashpot in parallel with an elastic spring of stiffness *E*_*i*_. The system exhibits two distinct timescales: a short timescale *τ*_1_ = 1.9 ± 0.3 s and a long relaxation time, *τ*_2_ = 37 ± 12 s. The fit provides an estimate for the effective elastic response of the bilayer in indentation *E*_*i*_ ∼ 1.6 MPa. This effective stiffness value is in good agreement with finite elements method simulations (inset of Fig. 5b), obtained by indenting a bilayer composed of a thin elastic film (cuticle with Young’s modulus *E*_*f*_ = 100 MPa and thickness 200 nm) deposited on an elastic substrate (epithelial cells with Young’s modulus *E* = 100 kPa).

In the nanoindentation experiment, the long relaxation time can be neglected at the scale of the experiment (see Methods), and the model is reduced to a simple Kelvin-Voigt model consisting of a dashpot in parallel to an elastic spring with a single characteristic time. Assuming Hertz contact, the strain of a viscoelastic Kelvin-Voigt material under indentation can be described by the equation 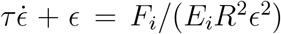, where *ϵ* = (*δ/R*)^1*/*2^ and *R* = 4.7 *μ*m the radius of the indenter. For a constant loading rate 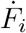, and in the limit of *t* ≪ *τ*, the strain can be approximated by *ϵ* ∼ (*t/T*)^2*/*3^, where 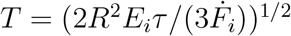. The experimental data collapse on this master curve (Fig. 5b), with a single value, *E*_*i*_*τ* being fitted to the curves. A similar procedure is applied to six samples, each subjected to a loading pattern of 21 loading and unloading cycles with increasing loading rates (see Fig. S6e). This yields *E*_*i*_*τ* ≈ 10 − 47 MPa.s and *τ* ≈ 6 − 29 s, in agreement with previous measurement of viscous time in epithelial cells [41]. FEM indentation of the bilayer, which accounts for the viscoelastic properties of the substrate with a viscous time scale *τ* = 10 s, as depicted by the dashed green curve in Fig. 5b, aligns with the experimental curves.

We extend our quasistatic model to the dynamic case by taking into account the viscoelasticity of the material. In this scenario, the hyperelastic material has an internal time scale *τ*, and is subjected to a pressure step 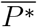 at *t* = 0. The model for *λ* as a function of *t/τ* (solid line in Fig. 5c) is supported by FEM simulations (dashed line Fig. 5c and Supplementary Movie 11). Snapshots of the FEM are shown at *t/τ* = 0, 5, and 25, illustrating how the system deforms both in-plane and vertically. The system ultimately reaches its equilibrium state within a characteristic time of ≈ 10*t/τ*. This prediction is compared with experimental measurements of the time evolution of *λ* obtained by bright-field microscopy. We plot *λ* as a function of time normalized by the measured internal timescale of the material obtained with the indentation experiment: *τ* = 10 s. The experimental data align with the theoretical prediction and the FEM without additional fitting parameters. The equilibrium state of the expansion and its dynamics are well captured by this model, in which a visco-hyperelastic material is subjected to an increase in pressure. This provides evidence that the expansion is solely due to an increase in pressure and that its dynamics are regulated by the internal timescale of the wing material itself.

## Discussion

Wing expansion in insects results from the coupling between material properties, geometry and fluid loading. Using the fruit fly *Drosophila melanogaster* as a model system, we first describe the complex structure of the folded wing, which consists of two sheets of epithelial cells separated by a lumen and covered by a wrinkled rigid cuticle. The sheets are held together by a hexagonal lattice of pillars containing microtubules. Expansion occurs a few minutes after emergence, as hemolymph flows in the lumen under a pressure increase of a few kPa. During expansion, we observe macroscopic unfolding of the wing, along with tissue stretching. This stretching is due to the unwrinkling and flattening of the volcano-like 3D microscopic structure of the apical surface of each cell. We then characterize the mechanical properties of the wings. Taking into account the geometry and strain stiffening hyperelasticity due to cuticle unwrinkling of microtubules stretching, we develop an effective model that analytically predicts in-plane expansion around an operating value of *P/E*. Finally, we extend our analysis to the dynamics of expansion. We model the viscoelastic behavior of epitelial cells using nanoindentation experiments. The implementation of dissipation in the effective model captures the expansion dynamics and is in agreement with experimental observations and finite element calculations.

Our model is based on an effective description of the complexity of the biological response and depends only on a finite set of parameters measured independently in the experiments: an elastic Young’s modulus *E*, a strain stiffness parameter *J*_*m*_ and a viscous time *τ* of the material. Despite its simplicity, the model gives a remarkable account of the kinematics and dynamics observed *in vivo*. In particular, it predicts the existence of an operating pressure *P/E* = 0.18 at which most of the expansion occurs. The pressure measured *in vivo* is slightly lower than the pressure predicted by the model or observed when we artificially inflate sacrificed flies after their removal from their pupal case. This discrepancy may be due to the fact that, for the sake of simplicity, we model the horizontal plates (cuticle-covered cells) and vertical pillars (microtubules) with the same effective constitutive law. Another potential source of discrepancy may be that tensile and indentation tests are carried out on dissected folded wings. Evaporation or cuticle hardening could artificially stiffen the moduli compared to *in vivo* conditions. Differential evaporation between dorsal and ventral plates could also explain the observation of the *Curly* phenotype when wings are artificially inflated in sacrificed flies.

We proposed unprecedented morphometric and mechanical characterization of the wing expansion. A physical model summarizes our results and shows that insects exploit the coupling between geometric and material non-linearities to achieve wing expansion around an operating point. This physical actuation is biologically triggered after eclosion by a hormone-controlled behaviour: the repeated intake of air and abdominal muscle contractions. This provides an interesting example of post-developmental morphogenesis. The groundwork is laid during larval and pupal development when the wings form, grow, undergo eversion and finally fold in the pupal case. However, upon eclosion the wings are not yet functional. The massive shape change that occurs as a result of air ingestion and abdominal contractions is mandatory to make them operational, and can be delayed by the insect if conditions are unfavorable [42]. Wing deployment is therefore a peculiar morphogenetic process: on the one hand, it takes place after development, and does not require cellular growth or mechanical activity, and on the other it requires active behavior of the animal to occur.

Our work paves the way for future studies aimed at understanding the pressure-driven expansion of deployable biological structures, and addressing the wide diversity of wing sizes and shapes in insects [43, 44]. In addition to its interest in fundamental biomechanics, improving our understanding of the mechanics of such deployable structures will find applications in the fields of material morphing physics and the engineering of flexible structures.

## Acknowledgments

We thank Marin Lebreuilly for help with initial indentation experiments, Nicolas Brouilly and the electron microscopy facility of the IBDM, Perrine Chaurand and the MATRIX plateform for the micro-tomography. This work was supported by Agence National de la Recherche (ANR Tremplin BioSoftAct), SH acknowledges funding from Fondation pour la Recherche MeÂdicale (FRM : FDT202304016556).

## Methods

### 1 Fly imaging

#### Strains and stock

Wild type Oregon-R flies are used throughout this study, with the exception of spinning disk fluorescent images, for which Ecad:GFP and Utrophin:GFP are used. Flies are maintained on standard fly food at 18°C in a temperature-controlled chamber with 12h light±dark cycles.

#### Bright-field microscopy

To record the expansion of the wings, a fly is collected immediately after eclosion from its pupal case and placed on a CO_2_ pad to anesthetize and manipulate it. A micro-manipulator is used to approach and glue a thin fiber (120 *μ*m diameter) to the dorsal part of the thorax of the fly, ensuring that the wings are free to move. The fly is then raised above the pad and the expansion of the wings is recorded from both a top view using a binocular microscope (Leica MZ16) and a lateral view using a Leica Z16 APO lens connected to two cameras (Imaging Source 37DFKBUX264).

#### Hemolymph flow vizualization

To visualize hemolymph flow through the wings during expansion, fluorescent beads are injected into the fly’s abdomen (500 nm pink beads, Drummond nanoject II). The fly is then attached to a thin stick and placed under a fluorescent binocular microscope (Nikon SMZ1000).

#### Profilometer characterization of the w rinkles

Wings are dissected and molded on uncured elastomer (Kerr Polyvinylsiloxane Impression Material type 3). The use of a mold provides a rigid sample after polymerization eliminating hairs that could mask surface features. Subsequently, the molds are scanned using a profilometer with an optical pen (micromesure 2, optical pen CL1-MG140, 1.3 *μ*m lateral resolution, 48 nm axial resolution). The surface 3D coordinates are analyzed using ImageJ and Matlab. Any overall surface curvature is corrected, the Voronoi tessellation of the wrinkles is generated, and the hexagonal organization, typical height, distance between wrinkles, and surface area of individual cells in both folded and expanded stages are extracted (Fig. 3c and yellow boxplot Fig. 3e for quantification).

#### Micro-CT

Micro-CT is performed at CEREGE (UM34, MATRIX platform). Flies are chemically fixed (glutaraldehyde + phosphate-buffered saline + paraformaldehyde), dehydrated with ethanol and dried with supercritical CO_2_ (critical point bypass) and placed in a kapton tube for imaging. A male is scanned over a 12-hour period, with the scan centered on the wings with a x20 objective (resolution of 0.8 *μ*m/voxel). A female is scanned over a 27-hour period, with the scan centered on a specific region of interest on the right wing with a x40 objective (0.32 *μ*m/voxel). ImageJ (version 2.14.0/1.54f) is employed for post-treatment, 3D image visualization (plugins 3D viewer and volume viewer), veins tracking, and geometrical characterization of the wings. The geometrical parameters (plates thickness *e*, pillars diameter *d*, pillars height *h*, and interpillar distance *a*) are measured by threshold, ultimate points and Voronoi tesselation processes on high-resolution scans (see Fig. 1b(ii-iii) and Fig. S2).

#### Transmitted Electron Microscope

Transmitted electron microscopy (TEM, FEI Tecnai g2 200 kv) is conducted at IBDM electronic microscopy platform. Wings are dissected from newly emerged flies (i) and adult fl ies 4h after wings expansion (ii). Wings are then fixed using cryofixation and cryosubstitution. The total wing thickness is quantififed before and after expansion using ImageJ (see boxplot wing thickness in Fig. 3e for quantification).

#### Spinning disk confocal microscope

A spinning disk confocal microscope is used to obtain the epithelial cells contour and the 3D apical cell surface from folded and expanded wings. The microscope is equipped with CSU-X1 spinning disk unit (Yokogawa) mounted on a Nikon Ti eclipse stand, 100x/1.49NA378 Nikon objective, EMCCD Andor iXon3 DU897, and MicroManager software. Folded wings are dissected from newly emerged flies, placed on a microscope slide, and covered with a drop of oil to avoid desiccation. A similar protocol is employed for expanded wings, which must be dissected shortly after wing expansion as epithelial cells undergo apoptosis and are washed out from the wing after expansion. Consequently, mature wings no longer exhibit fluorescence. The cell contours of Ecad:GFP flies are obtained with the Tissue Analyzer plugin in ImageJ, and quantification (cell surface area, cell-to-cell distance) is conducted in Matlab (MathWorks R2023b). The apical cell surface is imaged by performing a high resolution z-stack on Utrophin:GFP wings. Utrophin, an Actin-binding protein, tags the apical cell cortex. The scans are visualized and thresholded in ImageJ, and the 3D surface areas of individual cells are integrated in Matlab (see Supplementary Movie 8 and inset in Fig 3c).

#### Two-photon microscopy

A two-photon microscope, equipped with a Zeiss 510 NLO (Inverse - LSM), a femtosecond laser (Laser Mai Tai DeepSee HP) and a 40 x/1.2 C Apochromat objective, is used to measure the total wing thickness *in vivo*. The fly thorax i s attached to a s tick, and the insect is placed such that the wing plane is aligned with the optical path of the microscope. Wing expansion is concomitant with body movement on the order of the size of the insect (∼ mm), which makes the measurement of the wing thickness (∼ 10 *μ*m) during the expansion process in a live fly somewhat c hallenging. Nevertheless, we capture fast z-stacks of the wing section starting from the wing distal tip. The analysis is conducted using ImageJ (see Supplementary Movie 7 and Fig. 3e for quantification). We measure a total wing thickness (composed of the height of the pillars *h* and the thickness of the plates *e*) of ≈ 36 *μ*m during expansion i.e. a 2-fold increase compare to folded wings cross-section. This yields a stretch of the pillars in the vertical direction of *λ*^*p*^ ≈ 4.8.

### 2 Mechanical measurements

#### Pressure

The internal pressure of the insect is quantified as the wings expand using an in-house experimental setup, mounted on a vibration-isolated optical table. The pressure probe is composed of a pressure sensor (Honeywell 24PCBFA6D) connected to a rigid microfluidic channel. A glass capillary (Clark capillary glass GC100-15, 1 mm outside diameter, 0.58 mm internal diameter, tip size 10-50*μ*m) previously pulled (Sutter Instrument P97-4832) is filled with a low viscosity silicon (AS 4 Wacker-Chemie, 0.004 m Pa.s) and secured to one end of the channel, while a motorized syringe at the other end allows for volume control of the system. The tip of the syringe is filled with deionized water to create an oil/water meniscus in the large section of the c apillary. This prevents an oil/hemolymph interface at the capillary tip, which could result in an additional Laplace pressure. The position and the shape of the oil/water meniscus are tracked to correct for any additional pressure due to a possible flux (hydraulic resistance) or non-flat meniscus (Laplace pr essure). Flies with folded wings are collected immediately following their emergence from the pupal case and placed on the CO_2_ pad. A micromanipulator is utilized to puncture the insect scutellum with the capillary mounted on the pressure probe. The CO_2_ pad is then removed, and the wings expansion is observed from both top and side views. A LabVIEW interface (NI LabVIEW 2017) is employed to synchronize the pressure measurements, volume control of the syringe, and the images captured by the two cameras.

#### Tensile tests

Tensile tests are conducted on folded fly wings to characterize their mechanical properties. Wings are dissected from newly emerged flies and glued at their proximal extremity to a motorized linear-stage (glue Loctite AA 352, linear stage PI VT-80). The distal extremity of the wing is glued to a load sensor (Magtrol MBB-02-0.05) previously calibrated with known loads. A water tank is placed in close proximity to the wing to prevent desiccation. Following the application of the glue, a period of several minutes is allowed for it to harden before the linear stage is moved (see Supplementary Movie 9). The entire setup is controlled via a custom Labview interface, which includes monitoring the experiments with two cameras (providing top and side views), controlling the linear stage, and recording the force from the load sensor. The setup is placed on a vibration-isolated optical table. The raw measured force, *F*, is normalized by the cross-sectional surface area *S*_0_ of the folded wing obtained via micro-CT scan to extract the engineering stress *F/S*_0_ (see supplementary § Geometrical parameters measure from micro-CT scan for details on obtaining *S*_0_). Upon stretching, we assume incompressibility and the cross-sectional surface area of the wing to decrease as *S* = *S*_0_*/λ*. Consequently, the true stress *σ*, can be expressed as *σ* = *F/S* = *λF/S*_0_. We use ImageJ to track one of the longitudinal vein (underlined in green Fig. 4b and Fig. S3a) during stretching and compute the tissue stretch *λ* = *l/l*_*F*_ where *l* is the vein arclength and *l*_*F*_ is the corresponding initial folded length. The macroscopic stretch, Λ, is defined as the ratio between the end-to-end distance of the vein, *L*, and its corresponding initial value, *L*_*F*_. See Fig. 4b for a graph of *F/S*_0_ versus the macroscopic stretch Λ. The inset depicts the true stress, *σ*, as a function of *λ*. A linear fit at small stretch yields the Young’s modulus, *E*.

#### Nano-indentation

Nano-indentation experiments are conducted on folded wings using a nano-indenter (Hysitron TI Premier). The wings are dissected from flies just after emergence from the pupal case. The wing is then carefully placed on a thin layer of viscous epoxy glue coated on a microscope slide. This procedure allows the wing to slightly embed in the epoxy while leaving the wing top surface in the open air. Once the epoxy has cured, providing a rigid substrate for the wing, the microscope slide is secured in the nano-indenter with magnets. Two types of tests are conducted: (i) loading cycles with increasing loading rates (maximum force 5*μ*N, rates 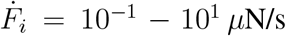) and (ii) creep tests with constant force (3 experiments at *F*_*i*_=3.2, 6 and 8.1 *μ*N). All experiments are performed within a maximum of 30 minutes after wing dissection. During this period, no time-dependent effects on the wing mechanical properties are observed. The raw force, *F*_*i*_, and indentation depth, *δ*, are analyzed using Matlab.

An indentation model of a viscoelastic material is constructed in order to extract the effective elasticity in indentation, *E*_*i*_, and the viscosity, *η*, of the wing. The loading cycles tests are well captured by a simple Kelvin-Voigt material undergoing Hertz-like indentation. The total stress of a Kelvin-Voigt material is the combination of a linear elastic response and a viscous dissipation, such that 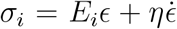. This model is then coupled with Hertz theory of contact *σ*_*i*_ = *F*_*i*_*/*(*Rδ*) and *ϵ* = (*δ/R*)^1*/*2^. The strain of a viscoelastic Kelvin-Voigt material under indentation is thus the solution of the equation:

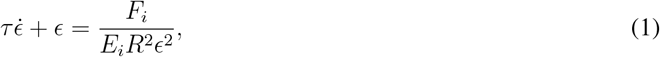

with *τ* = *η/E*_*i*_. Indentation cycles are performed at constant loading rate, so that 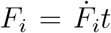, with 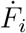 increasing with each cycle. In this case, strain can be solved from Eq. 1:

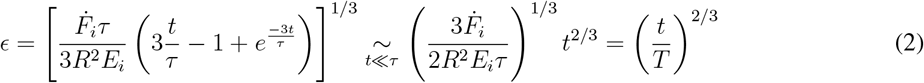

where 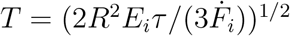. The experimental data are observed to collapse to a single line with a slope of 2/3 in a log-log scale in Fig. 5b. The data are fitted to the model in this limit to find *η* = *E*_*i*_*τ* ≈ 10 − 47 MPa.s. This range corresponds to the dispersion obtained by applying this procedure on six different samples, each subjected to a loading pattern of 21 loading and unloading cycles with an increasing loading rates.

The experimental creep test comprises two distinct phases: a loading phase during which the force is increased at a constant rate, and a subsequent phase in which the force remains constant (see the force curve in gray in the Inset of Fig. 5a). The resulting strain *ϵ* relaxed slowly over time, a behavior which is well captured by a three elements Maxwell-Jeffrey model comprising a dashpot *η*_2_ in series to a Kelvin-Voigt element (a dashpot *η*_1_ in parallel with a spring *E*_*i*_).

When coupled with Hertz contact, we obtain:

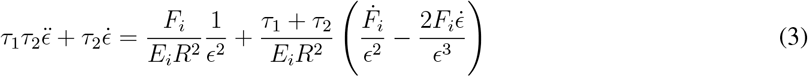

where *τ*_1_ = *η*_1_*/E*_*i*_ and *τ*_2_ = *η*_2_*/E*_*i*_. Eq. 3 is solved numerically, and the creep experimental data are fitted with *E*_*i*_, *τ*_1_ and *τ*_2_ as fitting parameters (see Inset Fig. 5a, yellow line, for the fit).

Note that in the nano-indentation experiments, the solutions of the three-element Maxwell-Jeffrey model reduce to the simpler Kelvin-Voigt model described above. A loading of the form 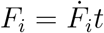 is imposed and we look for solutions of the form *ϵ* ≈ (*t/T*)^*α*^. For *α* = 2*/*3, there are two groups of terms that evolve as ∼ *t*^−1*/*3^ and ∼ *t*^−4*/*3^. For short times, the dynamics is governed by terms in power of −4*/*3. Balancing these terms, we find the expression *ϵ*(*t*) ≈ (*t/T*_0_)^2*/*3^, with

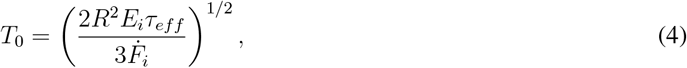

where *τ*_*eff*_ = *τ*_1_*τ*_2_*/*(*τ*_1_ + *τ*_2_) which tends toward *τ*_1_ for *τ*_1_ ≪ *τ*_2_. Numerical simulations of Eq. 3 for different values of 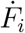 confirm the above scaling.

### 3 Model

#### Wing geometry and force balance

We consider two infinite plates of thickness *e*, connected with p illars of diameter *d*, height *h*, organized in a hexagonal lattice of pitch *a*. We assume that both plates undergo spatially homogeneous equibiaxial extension in e_*x*_ and e_*y*_, so that the principal stretches in the plates read *λ*_*x*_ = *λ*_*y*_ = *λ*. We assume that the material is incompressible, which implies that *λ*_*x*_*λ*_*y*_*λ*_*z*_ = 1 and *λ*_*z*_ = 1*/λ*^2^. For the pillars, we consider a uniaxial extension in the e_*z*_ direction. Incompressibility and isotropy lead to the following equation 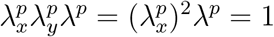, superscript *p* refers to the pillars and we denote 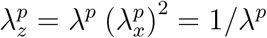.

A force balance in the plates relates the Cauchy stress in the principal directions *σ*_*XX*_ = *σ*_*Y Y*_ = *σ* to the internal pressure *P* [37]. As the plates become thinner by a factor of 1*/λ*^2^ and the pillar height increases by *λ*^*p*^, we obtain the equation 2 *σe/λ*^2^ = *Phλ*^*p*^ and:

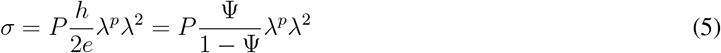

with Ψ = *h/*(*h* + 2*e*) the relative height of the pillars in the non-deformed geometry.

Similarly for the pillars, the longitudinal Cauchy stress 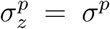 compensates for the increase in internal pressure, so that *σ*^*p*^*A*^*p*^*/λ*^*p*^ = *p* (*Aλ*^2^ − *A*^*p*^*/λ*^*p*^) where *A*^*p*^ is area of the pillars in the plane and *A* is the total area of the plates in the plane. We introduce the density of the pillars in the plane 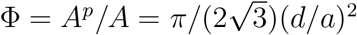, such that:

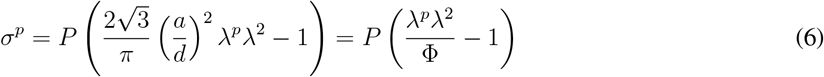

In order to solve these equations and obtain the values of (*λ, λ*^*p*^) as a function of *P*, it is necessary to define the material’s constitutive law.

#### Linear elastic model

In this section, we assume that the structure is composed of an isotropic homogeneous linear elastic material with Young’s modulus *E* and Poisson ratio *ν*. For the stress in the plane of the plates, the theory of linear elasticity yields:

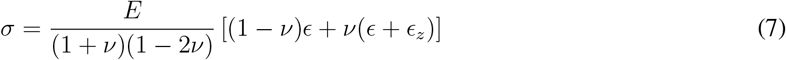

with the in-plane strain *ϵ* = *λ* − 1 and in the normal direction *ϵ*_*z*_ = *λ*_*z*_ − 1. We assume that the plates are under plane stress *σ*_*z*_ = 0, the thickness *e* being small compared to the other dimensions. We obtain (1 −*ν*)*ϵ*_*z*_ +2*νϵ* = 0, which gives *ϵ*_*z*_ = −2*νϵ/*(1 − *ν*), which we replace in Eq. 7 to obtain:

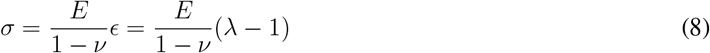

Similarly for the pillars, we write the general expression for the stress in the principal direction:

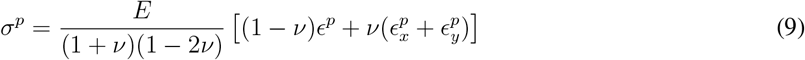

We explicit 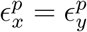 by writing the stress in one of the horizontal direction:

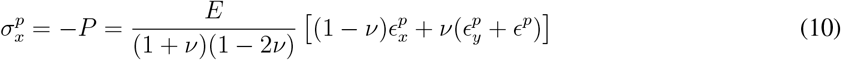

Indeed, the pillars are compressed by the internal pressure. We find 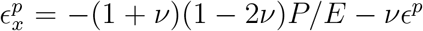, that we inject in Eq. 9 to get:

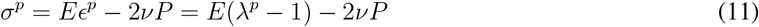

With Eq. 5-6, we obtain the following system of coupled non-linear equations:

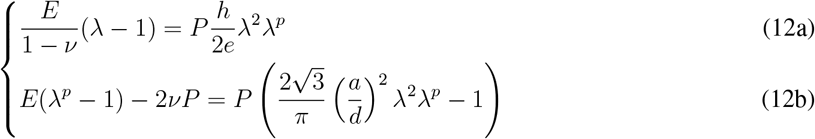

or equivalently with the non-dimensional pressure 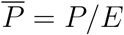:

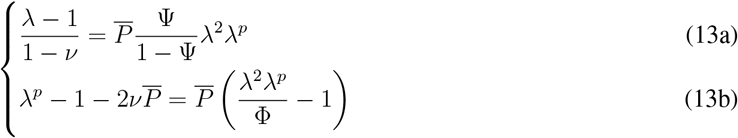

Eq. 13b gives 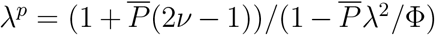 when 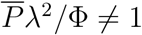. This result is then inserted into Eq. 13a to obtain the following equation for *λ*:

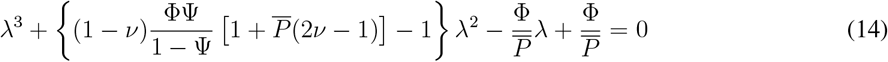

This linear elastic model predicts a finite pressure at which the stretch, both in the horizontal and vertical direction, diverges. This would be detrimental for the wing, which requires a large but finite expansion at a given pressure. Furthermore, this simple model does not take into account the microstructure of the wing: (i) the epithelial cells are covered by a rigid cuticle initially wrinkled that yields to a strain stiffening as the tissue expands; and (ii) the two bilayers are connected with microtubules that uncoil as the plates move apart until the stress in the pillars diverges when these filaments get straight. The aforementioned features, as illustrated Fig. 4a, prompt us to adopt a more realistic model that considers large strains and nonlinearities (strain stiffening) in the material.

#### Hyperelastic Gent model

We use the phenomenological Gent hyperelastic model to describe the strain stiffening. This constitutive model is characterized by two parameters: the shear modulus *μ* = *E/*(2(1 + *ν*)) and a limiting value *J*_*m*_ of the left Cauchy-Green deformation tensor first invariant. The second Piola-Kirchhoff stress tensor *S* depends on the strain energy density *W* :

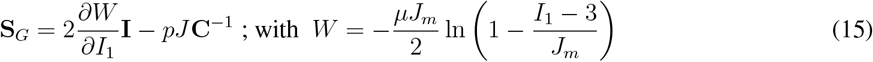

where **C** = **F**^*T*^ **F** is the right Cauchy-Green deformation tensor, **F** is the deformation gradient tensor, *I*_1_ = tr(**C**), *J* = det(**F**), **I** is the identity and *p* is a reactive pressure due to incompressibility constraint.

The plates are under plane stress *S*_*ZZ*_ = 0 yielding the in-plane stress:

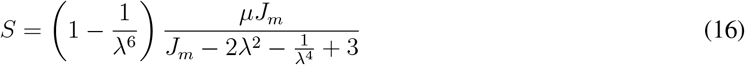

The pillars are compressed in the horizontal direction by internal pressure, 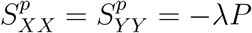, so that:

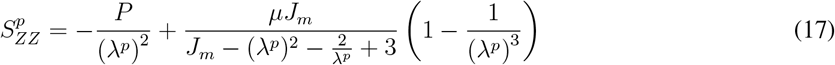

The Cauchy stress relates to the second Piola-Kirchhoff by ***σ***_*G*_ = *J*^−1^**FS**_*G*_**F**^*T*^ such that with Eq. 5-6, we obtain [37]:

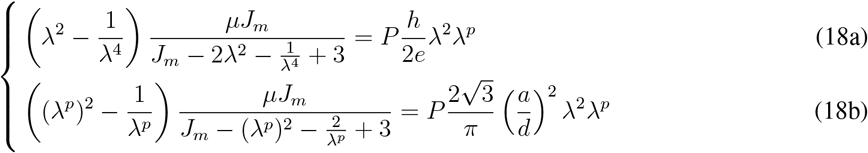

or in non-dimentionnal form with 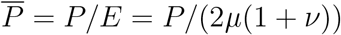:

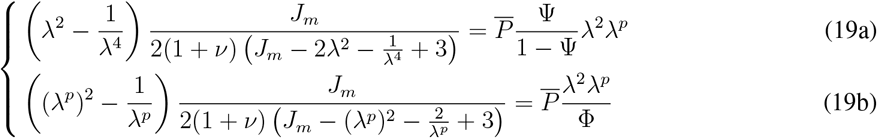

The deformations in the plane *λ* and in the pillars *λ*^*p*^ are predicted numerically by solving the coupled non linear equations as a function of the applied normalized pressure 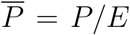 (see Fig. 4c, and Fig. S4 for a parametric study of the model).

#### Dynamical model

We take into account the viscous dissipation measured experimentally by extending the Kelvin-Voigt model to a hyperelastic material. The second Piola-Kirchhoff stress tensor in the viscous branch is 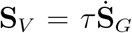 such that the total stress is 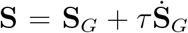. This yields ***σ*** = *J*^−1^***F SF*** ^*T*^ and, replacing the Cauchy stress in Eq. 5-6, we obtain:

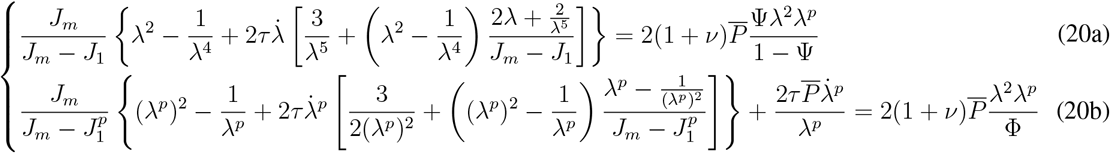

where *J*_1_ = *I*_1_ − 3 = 2*λ*^2^ + 1*/λ*^4^ − 3 and 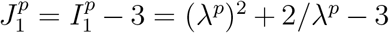. The expansion dynamics is predicted numerically by solving Eq. 20a-20b. In particular, Fig. 5c shows the in-plane deformation *λ* as a function of the normalized time *t/τ* for a constant applied pressure 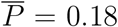. The experimental data (gray markers), for which the time is normalized by the measured viscous dissipation time *τ* = 10 s, exhibit a perfect match with the model, without any fitting parameters.

### 4. Finite Element Method

We perform finite element method simulations using the commercial software COMSOL Multiphysics (version 6.1) to (i) build a fundamental understanding of indentation experiments and (ii) support our model of wings expansion.

#### Indentation

We quantitatively study the effect of a bilayer on the indentation measurement by performing 2D axi-symmetric FEM simulations of a rigid spherical indenter (radius *R* = 4.7*μ*m, *E*_*ind*_ = 110 GPa, *ν*_*ind*_ = 0.3) applying a force to a bilayer composed of a soft cellular-like substrate (thickness 200*μ*m, width 400*μ*m) covered by a more rigid cuticle-like film 200 nm-thick. The substrate is modeled by a viscoelastic Kelvin-Voigt material with Young’s modulus *E* = 100 kPa and viscous time *τ* = 10 s, while the film is purely elastic with Young’s modulus *E*_*f*_ = 100 MPa. The boundaries are fixed at the bottom and the side of the system while loading-unloading cycles are applied on the bilayer by imposing the force with increasing loading rates.

#### Expansion

We numerically test our wing expansion model by conducting 3D FEM simulations of the pressure-driven deformation of a structure exhibiting the geometry of the wing: two plates with a thickness of *e* = 6.5 *μ*m connected with pillars of height *h* = 7.5 *μ*m, diameter *d* = 3.3 *μ*m, and an interpillar distance *a* = 6.2 *μ*m in a square lattice. *a* is chosen so that the in-plane pillars density in the FEM corresponds to that measured on micro-CT scans (see Fig.1b). The system is modeled using an isotropic and incompressible hyperelastic Gent material (Young’s modulus *E* = 100 kPa, limiting value *J*_*m*_ = 20) and solved for 1/8th of the geometry with symmetric boundary conditions. We first perform a stationary study, in which we impose an increasing pressure from *P* = 0 to 20 kPa and measure the in-plane deformation, *λ*, and vertical deformation, *λ*^*p*^ (see Fig. 4c black and orange dashed lines). We then turn to a time-dependent study by adding a viscous dissipation (Kelvin-Voigt viscous time *τ* = 10 s) in the material. We impose a constant pressure *P/E* = 0.18 at *t* = 0 and measure *λ* and *λ*^*p*^ over time. Snapshots of 1/2 of the geometry are shown at different times *t/τ* in Fig. 5c (see also Supplementary Movie 11).

## Supplementary information

### Macroscopic origami folding and vein network

Newly eclosed fly exhibits highly folded wings. Fig. S1a shows a folded fly wing (top: dorsal side; bottom: ventral side) with the veins used as landmarks throughout this study indicated in colors (see Fig. 2). This stereotypical folding, which occurs along the longitudinal veins but also in a perpendicular direction (ºmarginal foldº at the proximal end of the wing [1]), expands within minutes according to an equally stereotyped dynamic. Fig. S1b shows an adult wing and the associated vein network after the organ has undergone unfolding and tissue stretching.

### Geometrical parameters measure from micro-CT scan

We use ImageJ to extract the geometrical parameters of the folded wing from micro-CT scans. As sketched Fig.1b, the folded wing is composed of two plates of thickness *e* connected through pillars of diameter *d*, height *h* and organized in a hexagonal lattice of pitch *a*. We apply thresholds, ultimate points and Voronoi tesselation on high-resolution scans (see Fig. 1b(iii), 1 voxel = 0.32 *μ*m) to measure the diameter of the pillars *d* ≈ 3.3 *μ*m and the distance between them *a* ≈ 6.7 *μ*m. We take advantage of lower-magnification scans (Fig. 1b(ii) and Fig. S2a, 1 voxel = 0.8 *μ*m) to measure the plates thickness *e* and the gap between them *h*. We obtain a first binary mask by applying a threshold on the upper and lower plates (see Fig. S2b). The open surface area of this mask thus gives *S*_0_ = 2*el* where *l* is the arclength of the section. We then apply a closing process to obtain the full silhouette of the section (Fig. S2c). We obtain *l* by performing a skeleton process on this last mask and get the average plate thickness *e* = *S*_0_*/*(2*l*) ≈ 6.5 *μ*m. The close surface area of the mask also yields the surface *S* = (2*e* + *h*)*l*, which allows to calculate the average gap *h* = (*S* − *S*_0_)*/l* ≈ 7.5 *μ*m. The total thickness of the wing (composed of the plates thickness and the gap in between) while the wing is expanding is obtained with two-photon imaging (see Fig. S2d and Supplementary Movie 7)

**Figure 1.**
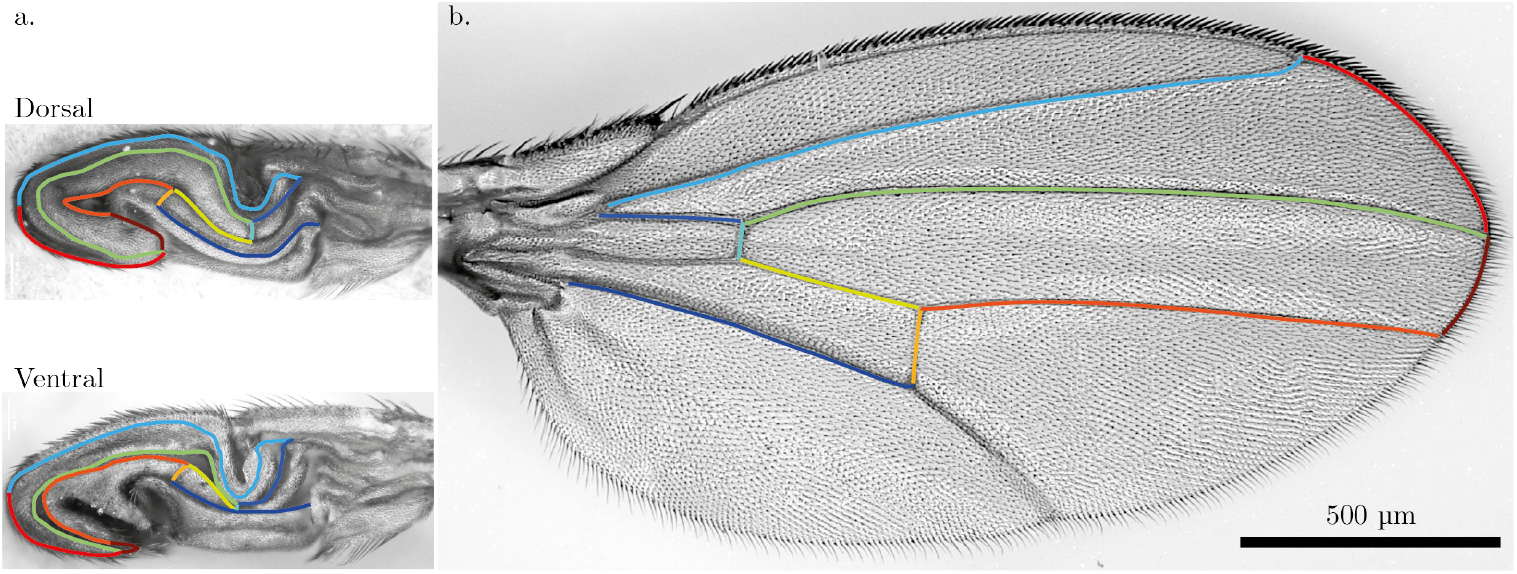
Folded and expanded wings with their corresponding vein network. **(a)** Dorsal and ventral side of a folded wing. Most of the veins follow the macroscopic folds and are located either on a mountain or in a valley fold. **(b)** Adult wing with its anterior edge at the top. The highlighted veins in a-b are the one that can also be visualized on the micro-CT scans.

**Figure 2.**
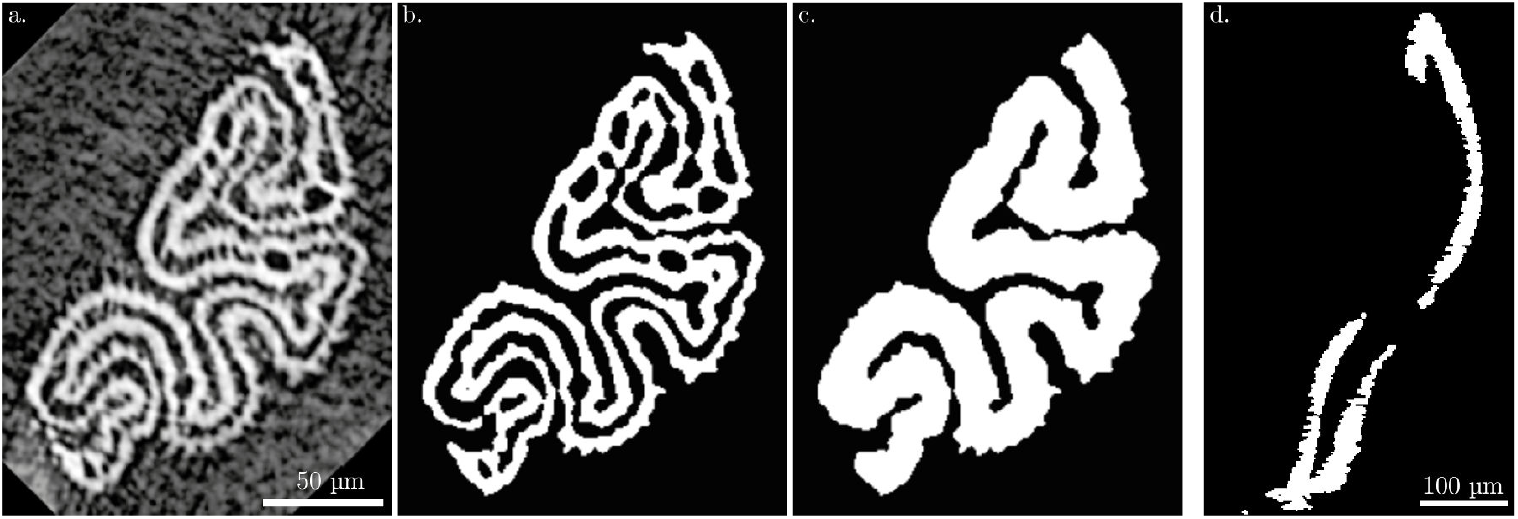
micro-CT and two-photon post-processing. **(a-c)** micro-CT: (a) Original cross-section normal to the proximo-distal axis of a folded wing. **(b)** Threshold highlighting the top and bottom plates. **(c)** A closing process yields the full silhouette of the section. (d) Two-photon: Cross section of an expanding wing.

### Wings mechanical properties measured through tensile tests

Tensile tests are conducted on dissected folded fly wings to characterize their mechanical properties. A displacement is imposed on the proximal end of the wing, which is attached to a linear stage. The force, *F*, is measured using a load sensor attached to the distal end of the wing (see picture of the experiment Fig S3a). Fig. S3b shows the stress for six experiments (colored markers) as a function of the stretch obtained by direct measurement of the variation in one of the longitudinal vein (shown in Fig S3a) and their associated linear fit at small strains (colored lines), which yield the Young’s modulus *E*. The average linear fit is shown as a dashed black line.

The strain stiffening of a wrinkled bilayer as it unwrinkles is illustrated using 2D FEM numerical simulations (COMSOL Multiphysics). The bilayer consists of a soft 10 *μ*m thick substrate of epithelial cells with a Young’s modulus *E* = 100 kPa covered by a 200 nm thick rigid film of cuticle with a Young’s modulus *E*_*f*_ = 100 MPa.

**Figure 3.**
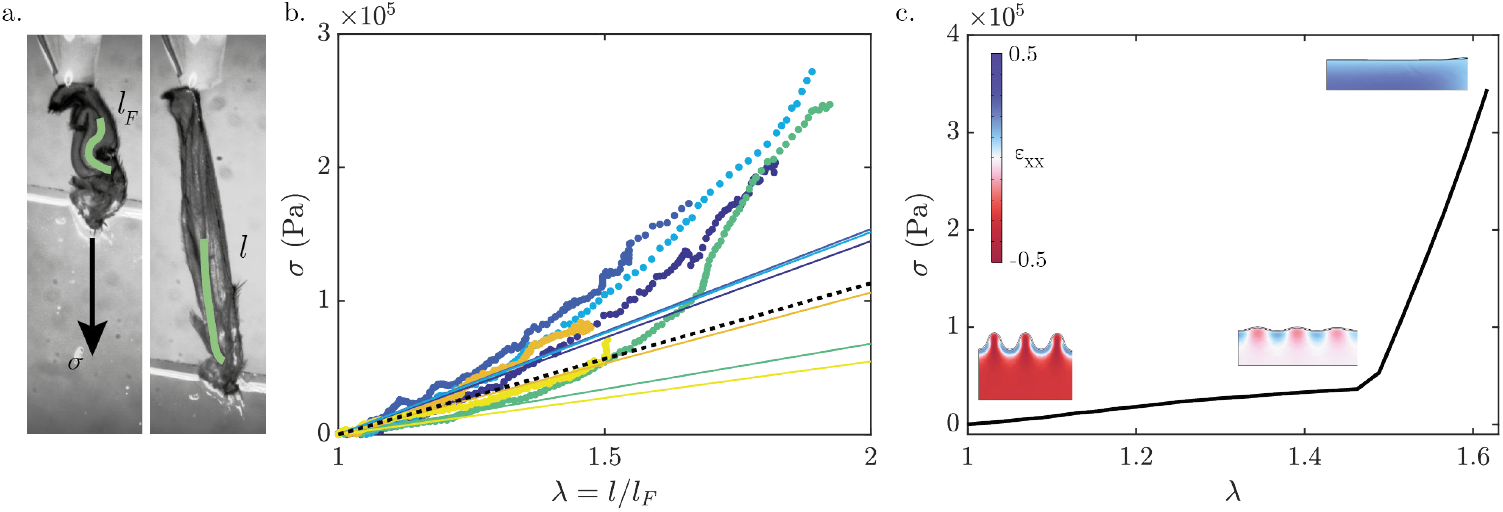
Tensile experiment. **(a)** Pictures of a tensile test experiment with the folded wing glued at both distal and proximal extremities. The followed vein is underlined in green. **(b)** Stress-stretch curve with all of the six experiments (data: colored markers; corresponding linear fit: colored solid lines; averaged fit: dashed black line). **(c)** FEM numerical simulation of the stretching of a wrinkled bilayer and computation of the associated stress-stretch. Color code is the strain in the tensile direction *ϵ*_*XX*_.

To improve convergence, we consider a hyperelastic Gent model for the substrate with an arbitrarily large value of *J*_*m*_ = 100, which has a negligible impact for the values of stretching considered in the simulation. We impose symmetric boundary conditions on the bottom and left sides of the system while we prescribe the displacement on the right side. The top surface is free. The initial state is obtained by thermally contracting the substrate to form wrinkles (see FEM snapshot, bottom left in Fig. S3c). We then impose a displacement to the right boundary. Fig. S3c shows the normal stress along the right boundary as a function of the stretch, *λ*. The curve exhibits two distinct regimes: (i) at moderate stretch (*λ <* 1.5), stress is low and is mainly dominated by the stretching of the soft substrate, while the unwrinkling of the rigid film has a minimal impact on the overall stiffness of the bilayer; (ii) as the wrinkles disappear (intermediate snapshot *λ* ∼ 1.5), the system becomes stiffer and the mechanical response is dominated by the stretching of the rigid film. This simulations illustrate the overall strain stiffening of the wing observed in the experiments, which stems from microscopic unwrinkling of the cuticle, and support the choice of the Gent’s hyperelastic model to account for the effective mechanical response of the composite tissue of the wing.

#### Model parameters

The wing expansion model depends on both mechanical parameters (Young’s modulus *E*, Gent limiting value *J*_*m*_) and geometrical features (plates thickness *e*, pillars height *h*, diameter *d*, interpillar distance *a*). The afore-mentioned parameters are quantified through microscopic characterization and mechanical testing. However, it is of interest to assess the impact of these parameters on model predictions, and the robustness of predictions to small parameter variations.

Fig. S4a illustrates that at small strain *λ <* 1.5, the linear elastic law does not differ greatly from a hyperelastic material. A purely linear elastic model predicts a finite pressure at which stretching, in both horizontal and vertical directions, diverges. We therefore proceed to a more realistic model that incorporates the effects of large strains and non-linearities (strain-stiffening) in the material. The value of *J*_*m*_ is determined by matching predictions of the strain in the pillars *λ*^*p*^ (Fig. S4b) with experimental observations. The maximum pillar strain measured *in vivo* is *λ*^*p*^ ∼ 4.8. A value of *J*_*m*_ = 20 corresponds well to this limiting value at working pressure 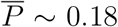. To keep the model simple, we have chosen to use the same parameter *J*_*m*_ for the biaxial expansion of the plates.

**Figure 4.**
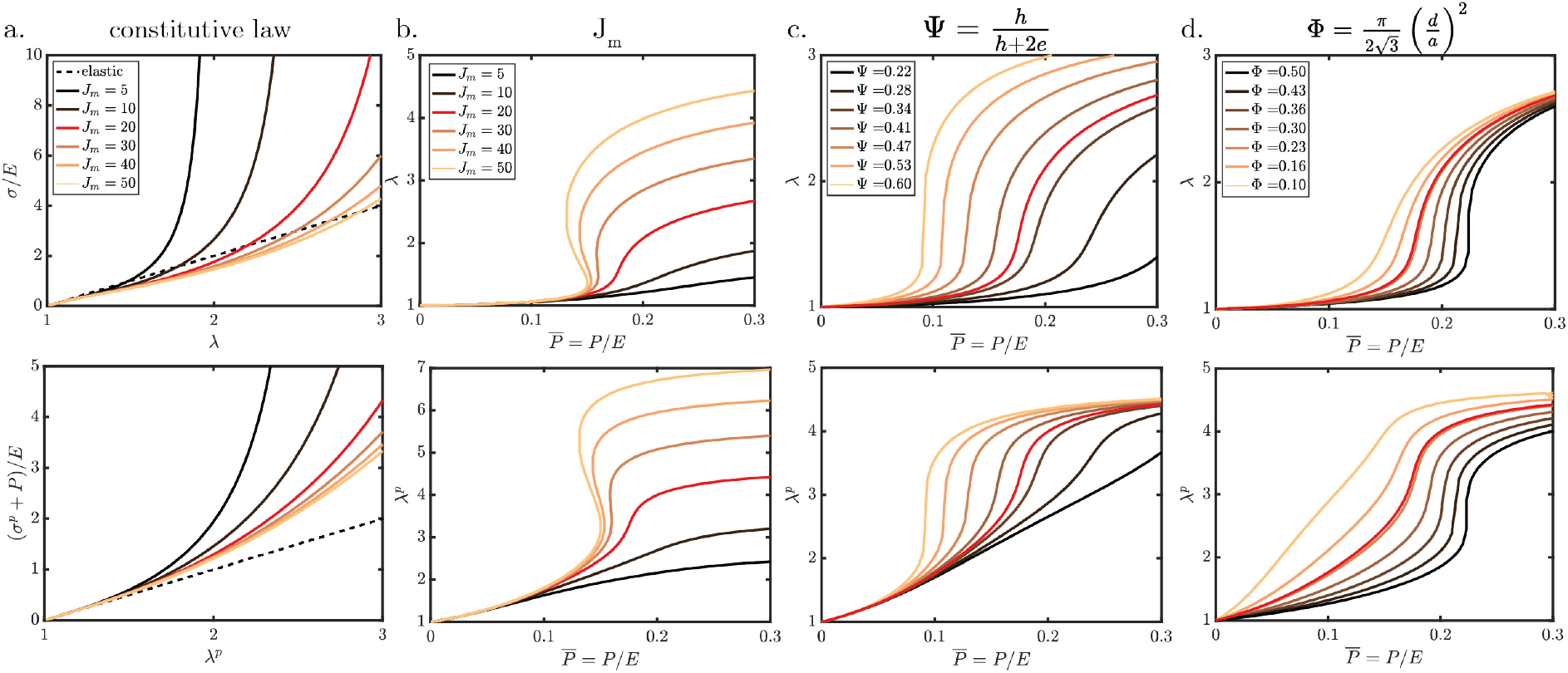
Model and influence of the mechanical and geometric parameters. **(a)** Elastic vs hyperelastic constitutive law: (top) in-plane normalized stress *σ/E* as a function of the in-plane stretch *λ* and (bottom) perpendicular normalized stress (*σ*^*p*^ + *P*)*/E* as a function of the pillar stretch *λ*^*p*^ for an elastic constitutive law (dotted line) and hyperlelastic Gent models (solid lines, different parameters *J*_*m*_). **(b-d)** (top) in plane stretch *λ* and (bottom) perpendicular stretch *λ*^*p*^ versus normalized pressure 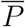 varying parameters *J*_*m*_ (b), pillars relative height Ψ (c), and in-plane pillars density Φ (d).

Figures S4c-d illustrate the influence of geometry on the prediction for the stretch in the plates *λ* (top graphs) and the stretch in the pillars in *λ*^*p*^ (bottom). We note in particular that more slender pillars (i.e. Ψ ∼ 1 or Φ ∼ 0 in terms of non-dimensional parameters) result in a shift of the curves towards lower working pressure.

To validate our model, we perform FEM numerical simulations of the inflation of a wing-shaped structure composed of a hyperelastic Gent material (*E* = 100 kPa, *J*_*m*_ = 20). Fig. S5a shows simulation predictions of the in-plane deformation *λ* as a function of the normalized applied pressure 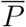 (dotted lines) for which we vary the system size (36, 64 and 37 pillars) and pillar arrangement (square and hexagonal lattice, adjusting the distance *a* between the pillars to keep the pillars density Φ = 0.22 constant, see FEM snapshots Fig. S5b). All predicted *λ* follow the same curve, which shows that (i) the system is sufficiently large for boundary effects to be negligible; and (ii) confirms that pillar organization has no key role in the model, the important parameter being the pillar density Φ. It should be noted that the model (shown in black solid line Fig. S5a) does not take into account the actual connection of the pillars to the membrane, which explains the discrepancy observed with the FEM.

**Figure 5.**
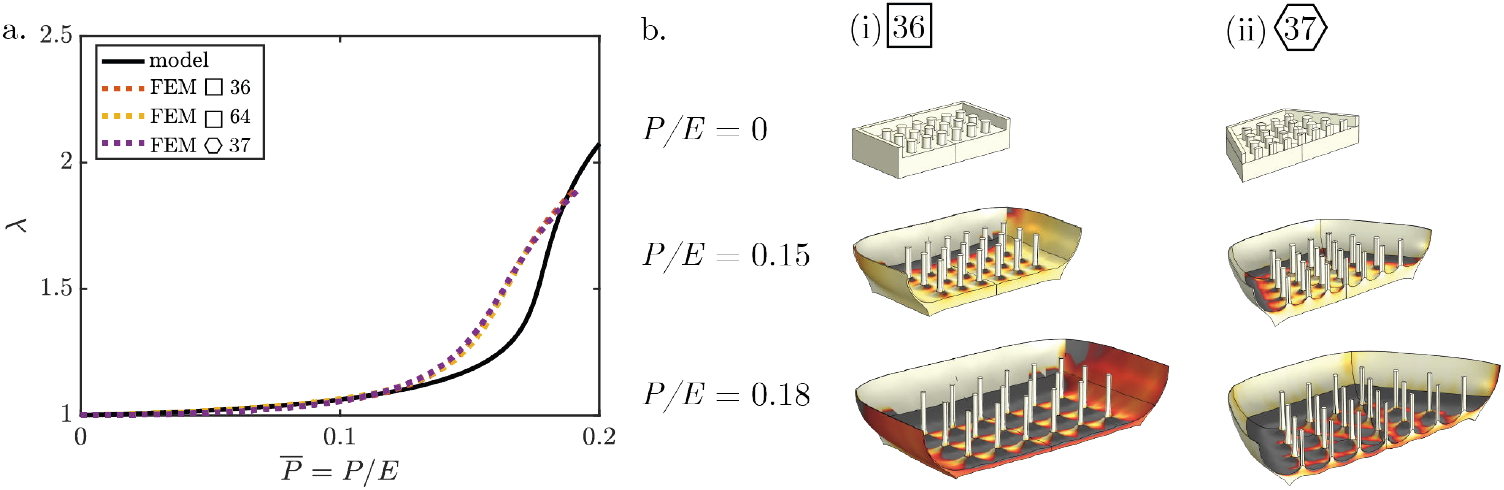
FEM simulations of inflating a hyperelastic wing-shaped structure. **(a)** Deformation *λ* as a function of the normalized applied pressure 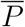 : all FEM predictions (dashed lines) varying the size of the system (36, 64 and 37 pillars) and the pillars organization (square and hexagonal lattice) collapse on the same curve. The model is shown as the black solid line. **(b)** FEM snapshots at different pressure of 1/4th of the geometry for (i) 36 pillars organized in a square lattice, and (ii) 37 pillars organized in a hexagonal lattice. Color code: strain in one of the in-plane principal direction.

#### Viscous dissipation probed through nanoindentation experiments

We perform creep tests and loading-unloading cycles in nanoindentation to measure the wing material viscous dissipation. Fig. S6a-c show three creep tests for which a constant force *F*_*i*_ is applied (see Fig. S6d for the loading signal). Each curve is fitted by a three elements Maxwell-Jeffrey model (yellow line), yielding to a short timescale *τ*_1_ = 1.9±0.3 s, a long relaxation time, *τ*_2_ = 37±12 s and an effective elastic response of the bilayer in indentation *E*_*i*_ ∼ 1.6 MPa.

Loading-unloading cycles tests in nanoindentation ± for which the long relaxation time can be neglected at the time scale of the experiments ± are well captured by a Kelvin-Voigt model as shown Fig. S6e. The experimental data (represented with colored markers, using 10 bins per sample to average all data series) follow the model *ϵ* = (*t/T*)^2*/*3^, where 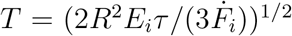, with a single value, *E*_*i*_*τ* being fitted to the curves. To confirm the model, we perform FEM numerical simulations of indenting a bilayer composed of a thin elastic film of Young’s modulus *E*_*f*_ = 100 MPa deposited onto a soft viscoelastic substrate (*E* = 100 kPa, *τ* = 10 s) at different loading rates (10^−1^ to 10^0^ *μ*N/s). The simulations, shown as dashed line Fig. S6e, also follow the master curve.

### *in vivo* pressure recording and artificial pressure increase

Fig. S7 shows the temporal evolution of pressure *P* (black curve) and deformation *λ* (red) for four distinct flies. We observe that pressure initially increases, reaching a plateau of ≈ 1.5 − 3.5 kPa during which most ± at least 50% ± of the wings expansion occurs.

We wonder whether wing expansion can be triggered by an external artificial increase of pressure. To test this hypothesis, a wild-type fly is placed under ether vapor for 10 minutes immediately after emerging from the pupal case. We then puncture its scutellum using a glass capillary (50-75 *μ*m outside diameter) connected to a syringe pump and a pressure sensor. PBS is injected to impose different pressure plateaus and the fly is observed for at least 2 minutes to identify any sign of expansion before increasing the pressure. No expansion of the wings is observed below the pressure plateau of *P* ∼ 10 kPa. Between *P* = 10 − 16 kPa, the wings expand in about 20 minutes and exhibit curly wings. Fig. S8a and Supplementary Movie 10 show top- and side-view snapshots of such an experiment. At higher pressure (*P >* 17 kPa), the pillars break, the ventral and dorsal layers delaminate, resulting in blisters or even balloon like wings (see Fig. S8b). Note that at this stage, increasing the pressure does not further stretch the tissue, which further justifies the use of a hyperelastic strain-stiffening model to describe the wing.

**Figure 6.**
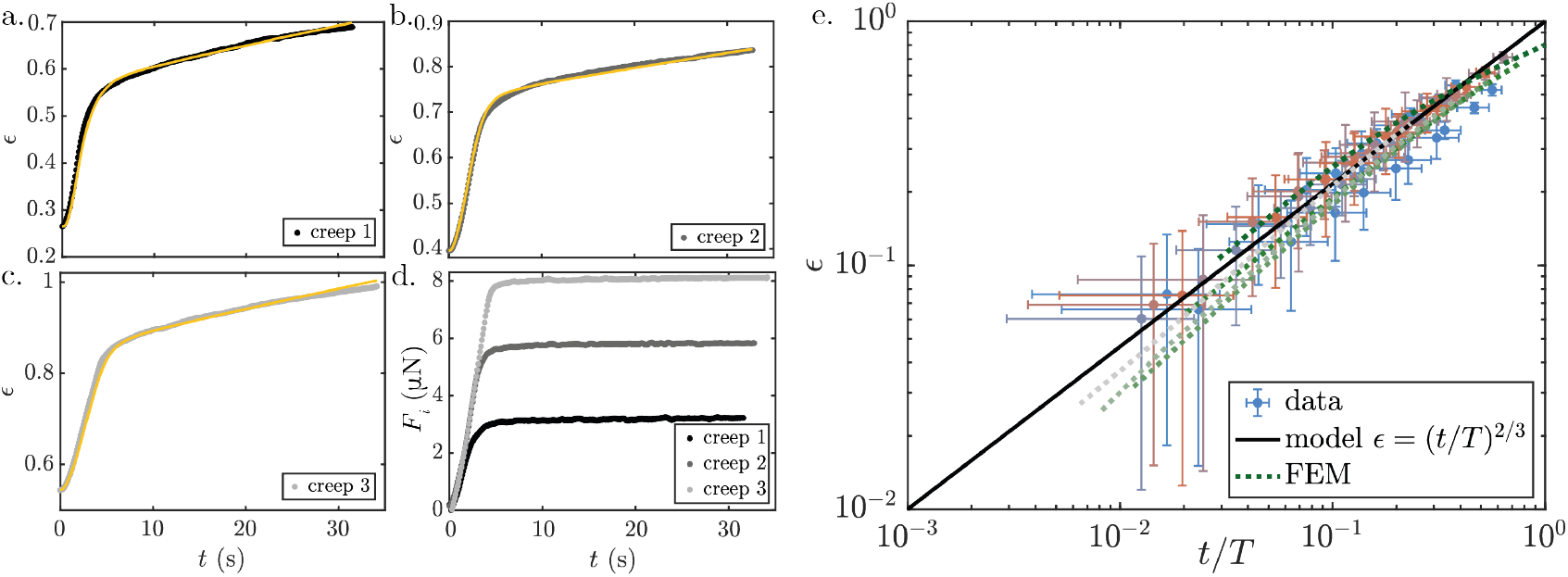
Nanoindentation on folded wings. **(a-c)** Deformation *ϵ* = (*δ/R*)^1*/*2^ as a function of time for three different creep experiments (experimental data: in black; Maxwell-Jeffrey fit: yellow line), each one undergoing a different force plateau *F*_*i*_ (see **(d)** for the corresponding loading signals). **(e)** *ϵ* versus normalized time *t/T* from nanoindentation experiments. Data (colored markers, using 10 bins per sample to average all experiments, standard deviation shown with errorbars) and FEM (dashed lines, from slow indentation 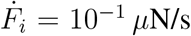 in dark green, to faster 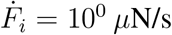 in gray) collapse on the model *ϵ* = (*t/T*)^2*/*3^ (black line).

**Figure 7.**
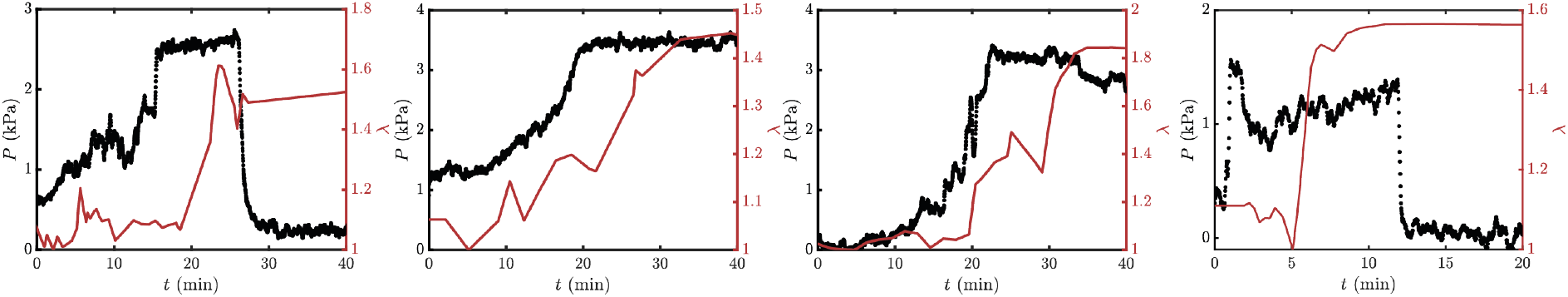
*in vivo* pressure measurements. Pressure recording (black curves) in 4 different flies and corresponding elongation *λ* (red) in time as the wings expand. Most of the wing expansion happens on a pressure plateau of 1.5 − 3.5 kPa.

**Figure 8.**
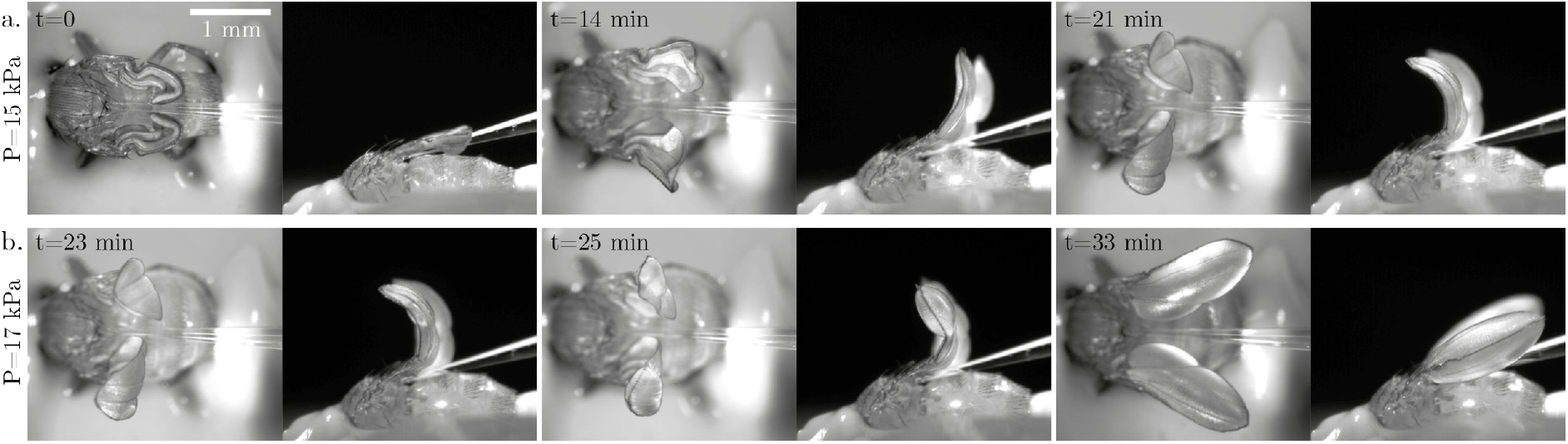
Artificial pressure increase. **(a)** A pressure plateau of *P* = 15 kPa is applied in a fixed wild type fly with folded wings, triggering curved wings expansion in ∼ 10 minutes. **(b)** An additional increase in the pressure to *P* = 17 kPa leads to microtubule pillars breakage and wing blade delamination resulting in blister or ballon-like wings.

## Description of supplementary movies

**Movie S1**. Wings expansion recording (wild type female *Drosophila*) from a top- and a side-view (timer mm:ss).

**Movie S2**. Side-view video of a fly increasing its internal pressure through air swallowing (white arrow points to pharyngeal pumping) and abdominal muscle contraction.

**Movie S3**. Micro-CT scan of a folded wing (ImageJ plugin 3D viewer). Z-stack of this same wing cross sections normal to the proximo-distal axis reveals the internal structure (namely the dorsal and ventral plates connected through pillars).

**Movie S4**. Fluorescent binocular microscopy of hemolymph flux marked with fluorescent beads during expansion (left) and in an adult wing (right). Hemolymph spreads the entire structure during expansion, that is highly different from hemolymph flows in an adult wing where it is only constraint to the vein network.

**Movie S5**. *in vivo* internal pressure measurement during wing expansion. We poke the scutellum of a newly emerged fly with a glass capillary connected to a pressure sensor and record wing expansion. Inset: measured pressure *P* (*t*) in time.

**Movie S6**. Combination of the recordings of wings expansion in 6 individuals shows reproducibility of the process (3 females on top, 3 males at the bottom). The time at which the wings start to unfold is chosen as a reference to combine the videos.

**Movie S7**. 3D reconstruction of the distal extremity of a wing during its expansion obtained with a two-photon microscope, allowing a measurement of the wing thickness *in vivo* during expansion of ∼ 35 *μ*m (to be compared with the ∼ 18 *μ*m thick wing before expansion, see TEM Fig.3b and quantification boxplot ºwing thicknessº Fig.3e).

**Movie S8**. 3D reconstruction of an individual cell apical surface (Utrophin:GFP) from z-stack obtained through fluorescent spinning disk microscopy. Integration of this 3D shape leads to a measurement of individual apical cell surface area (see Folded condition of the boxplot º3D Areaº, Fig. 3e).

**Movie S9**. Tensile test performed on a dissected folded wing. The distal tip of the wing is connected to a force sensor while we impose a displacement to the proximal extremity glued to a moving stage (see Fig. 4b and Fig. S3a for the measured stress-strain curves).

**Movie S10**. An artificial pressure increase triggers wings expansion. We poke the scutellum of a fixed newly emerged wild type fly with a glass capillary connected to a syringe and impose a first pressure plateau of 15 kPa for which wings unfold and curl upward. At *t* = 23 minutes an additional increase in the pressure plateau to 17 kPa leads to microtubule pillars breaking and delamination of the dorsal and ventral layer resulting in balloon-like wings (see Fig. S8 for corresponding snapshots).

**Movie S11**. 3D FEM simulations (COMSOL Multiphysics) of the inflation of wing-like structure made of two plates of thickness *e* = 6.5 *μ*m connected with pillars (height *h* = 7.5 *μ*m, diameter *d* = 3.3 *μ*m, interpillar distance *a* = 6.2 *μ*m). The visco-hyperlastic material (*E* = 100 kPa, *J*_*m*_ = 20, *τ* = 10 s) is submitted to a pressure step of *P* = 16 kPa at *t* = 0. Color bar: in-plain strain.

## References

[1] Nelson, C. M. On buckling morphogenesis. J. Biomech. Eng. 138, 021005 (2016).

[2] Shyer, A. E. et al. Villification: how the gut gets its villi. Science 342, 212±218 (2013).

[3] Tallinen, T. et al. On the growth and form of cortical convolutions. Nat. Phys 12, 588±593 (2016).

[4] Kim, S., Pochitaloff, M., Stooke-Vaughan, G. A. & Campxàs, O. Embryonic tissues as active foams. Nat. Phys 17, 859±866 (2021).

[5] Mitchell, N. P. et al. Visceral organ morphogenesis via calcium-patterned muscle constrictions. Elife 11, e77355 (2022).

[6] Etournay, R. et al. Interplay of cell dynamics and epithelial tension during morphogenesis of the drosophila pupal wing. Elife 4, e07090 (2015).

[7] de la Loza, M. D. & Thompson, B. Forces shaping the drosophila wing. Mech. Dev. 144, 23±32 (2017).

[8] Harmansa, S., Erlich, A., Eloy, C., Zurlo, G. & Lecuit, T. Growth anisotropy of the extracellular matrix shapes a developing organ. Nat. Commun 14, 1220 (2023).

[9] Tsuboi, A., Fujimoto, K. & Kondo, T. Spatiotemporal remodeling of extracellular matrix orients epithelial sheet folding. Sci. Adv. 9, eadh2154 (2023).

[10] Eidmann, H. Untersuchungen uÈber wachstum und haÈutung der insekten. Z. Morph. OÈ kol. Tiere 2, 567±610 (1924).

[11] Cottrell, C. The imaginal ecdysis of blowflies. observations on the hydrostatic mechanisms involved in digging and expansion. J. Exp. Biol. 39, 431±448 (1962).

[12] Moreau, R. Variations de la pression interne au cours de l’eÂmergence et de l’expansion des ailes chez bombyx mori et pieris brassicae. J. Insect Physiol. 20, 1475±1480 (1974).

[13] Dewey, E. M. et al. Identification of the gene encoding bursicon, an insect neuropeptide responsible for cuticle sclerotization and wing spreading. Curr. Biol. 14, 1208±1213 (2004).

[14] Honegger, H.-W., Dewey, E. M. & Ewer, J. Bursicon, the tanning hormone of insects: recent advances following the discovery of its molecular identity. J. Comp. Physiol. A 194, 989±1005 (2008).

[15] Pellegrino, S. Deployable structures in engineering. In Deployable structures, 1±35 (Springer, 2001).

[16] Klein, Y., Efrati, E. & Sharon, E. Shaping of elastic sheets by prescription of non-euclidean metrics. Science 315, 1116±1120 (2007).

[17] Silverberg, J. L. et al. Using origami design principles to fold reprogrammable mechanical metamaterials. Science 345, 647±650 (2014).

[18] Filipov, E. T., Tachi, T. & Paulino, G. H. Origami tubes assembled into stiff, yet reconfigurable structures and metamaterials. Proc. Natl. Acad. Sci. U.S.A. 112, 12321±12326 (2015).

[19] Dudte, L. H., Vouga, E., Tachi, T. & Mahadevan, L. Programming curvature using origami tessellations. Nat. Mater. 15, 583±588 (2016).

[20] Faber, J. A., Arrieta, A. F. & Studart, A. R. Bioinspired spring origami. Science 359, 1386±1391 (2018).

[21] Melancon, D., Gorissen, B., GarcÂia-Mora, C. J., Hoberman, C. & Bertoldi, K. Multistable inflatable origami structures at the metre scale. Nature 592, 545±550 (2021).

[22] SieÂfert, E., Reyssat, E., Bico, J. & Roman, B. Bio-inspired pneumatic shape-morphing elastomers. Nat. Mater. 18, 24±28 (2019).

[23] Kim, W. et al. Bioinspired dual-morphing stretchable origami. Sci. Robot. 4, eaay3493 (2019).

[24] Jones, T. J., Jambon-Puillet, E., Marthelot, J. & Brun, P.-T. Bubble casting soft robotics. Nature 599, 229±233 (2021).

[25] Jones, T. J., Dupuis, T., Jambon-Puillet, E., Marthelot, J. & Brun, P.-T. Soft deployable structures via core-shell inflatables. Phys. Rev. Lett. 130, 128201 (2023).

[26] Peabody, N. C. et al. Bursicon functions within the drosophila cns to modulate wing expansion behavior, hormone secretion, and cell death. J. Neurosci. 28, 14379±14391 (2008).

[27] White, B. H. & Ewer, J. Neural and hormonal control of postecdysial behaviors in insects. Annu. Rev. Entomol. 59, 363±381 (2014).

[28] Pass, G. Beyond aerodynamics: The critical roles of the circulatory and tracheal systems in maintaining insect wing functionality. Arthropod Struct. Dev. 47, 391±407 (2018).

[29] Ewer, J. & Reynolds, S. Neuropeptide control of molting in insects. In Hormones, brain and behavior, 1±XVI (Elsevier, 2002).

[30] Denlinger, D. L. & ZdaÂrek, J. Metamorphosis behavior of flies. Annu. Rev. Entomol. 39, 243±266 (1994).

[31] Barker, E. D. Inflatable mattress. United States Patent (1951).

[32] Saito, K., Nomura, S., Yamamoto, S., Niiyama, R. & Okabe, Y. Investigation of hindwing folding in ladybird beetles by artificial elytron transplantation and microcomputed tomography. Proc. Natl. Acad. Sci. U.S.A. 114, 5624±5628 (2017).

[33] Kiger Jr, J. A. et al. Tissue remodeling during maturation of the drosophila wing. Dev. Biol. 301, 178±191 (2007).

[34] ToÈgel, M., Pass, G. & Paululat, A. The drosophila wing hearts originate from pericardial cells and are essential for wing maturation. Dev. Biol. 318, 29±37 (2008).

[35] Johnson, S. A. & Milner, M. J. The final stages of wing development in drosophila melanogaster. Tissue cell 19, 505±513 (1987).

[36] BoÈkel, C., Prokop, A. & Brown, N. H. Papillote and piopio: Drosophila zp-domain proteins required for cell adhesion to the apical extracellular matrix and microtubule organization. J. Cell Sci. 118, 633±642 (2005).

[37] SieÂfert, E. & Roman, B. Morphogenesis through elastic phase separation in a pneumatic surface. C. R. Mec. 348, 649±657 (2020).

[38] Kenny, M. C., Giarra, M. N., Granata, E. & Socha, J. J. How temperature influences the viscosity of hornworm hemolymph. J. Exp. Biol. 221, jeb186338 (2018).

[39] Lechantre, A. et al. Microrheology of haemolymph plasma of the bumblebee bombus terrestris. J. Exp. Biol. 226, jeb245894 (2023).

[40] Elbaz, S. B. & Gat, A. D. Dynamics of viscous liquid within a closed elastic cylinder subject to external forces with application to soft robotics. J. Fluid Mech. 758, 221±237 (2014).

[41] Bambardekar, K., CleÂment, R., Blanc, O., Chardès, C. & Lenne, P.-F. Direct laser manipulation reveals the mechanics of cell contacts in vivo. Proc. Natl. Acad. Sci. U.S.A. 112, 1416±1421 (2015).

[42] Peabody, N. C. et al. Characterization of the decision network for wing expansion in drosophila using targeted expression of the trpm8 channel. J Neurosci. 29, 3343±3353 (2009).

[43] Salcedo, M. K., Hoffmann, J., Donoughe, S. & Mahadevan, L. Computational analysis of size, shape and structure of insect wings. Biol. Open 8, bio040774 (2019).

[44] Salcedo, M. K., Jung, S. & Combes, S. A. Autonomous expansion of grasshopper wings reveals external forces contribute to final adult wing shape. Integr. Comp. Biol. 63, 1111±1126 (2023).

## References

[1] Alice Tsuboi, Koichi Fujimoto, and Takefumi Kondo. Spatiotemporal remodeling of extracellular matrix orients epithelial sheet folding. Sci. Adv., 9(35):eadh 2154, 2023.

